# A TLS11a-decorated ionizable lipid nanoparticle platform and a multilevel-validated CRISPR LDLR-knockout HepG2 model for hepatocyte-preferential mRNA delivery

**DOI:** 10.64898/2026.07.21.739738

**Authors:** Irfan Hussain, Naji Kholaif, Raghad Alsultan, Rahma Eltahir, Huda Alajlan, Tanveer Ahmad Mir, Jahan Salma, Fawad Ur Rehman, Anas Mohammed Alazami, Farhatullah Syed

## Abstract

Ionizable lipid nanoparticles (LNPs) are widely used for delivery of CRISPR/Cas9 payloads to hepatocytes, but conventional hepatic uptake is strongly influenced by adsorption of apolipoprotein E and subsequent low-density lipoprotein receptor (LDLR)-mediated internalization. This dependence may limit specificity and reduce efficacy in LDLR-deficient settings. Here, we designed an aptamer-functionalized LNP platform to enable hepatocyte-selective genome editing through an LDLR-independent route and validated its performance using a genetically defined LDLR-knockout HepG2 model. Ionizable LNPs co-encapsulating Cas9 mRNA and an LDLR-targeting guide RNA were surface-decorated with the hepatocellular carcinoma-targeting TLS11a aptamer using thiol-maleimide chemistry. Comprehensive physicochemical analysis using cryo-electron microscopy, dynamic light scattering, pKa titration, UV and circular dichroism spectroscopy, X-ray photoelectron spectroscopy, and molecular beacon assays confirmed uniform nanoparticles of approximately 105 nm, preserved mRNA integrity, retained endosomal charge-switching behavior with a pKa of approximately 6.3 to 6.5, and maintained correctly folded surface-displayed TLS11a. TLS11a decoration increased Cas9 mRNA delivery to HepG2 cells from 39% to 79% Cy5-positive cells, while reducing uptake in receptor-low control cells, supporting aptamer-associated and cell-preferential delivery. In parallel, CRISPR/Cas9-mediated deletion of LDLR exon 2 generated a validated LDLR-deficient HepG2 line, confirmed at genomic, transcript, and protein levels. LDLR loss reduced LDL binding and uptake by approximately 85%, while transferrin uptake was preserved, indicating selective impairment of LDLR-dependent endocytosis. Cholesterol depletion activated the SCAP-SREBP-2 pathway and induced cholesterol biosynthesis genes. Together, these findings establish a modular aptamer-guided LNP system for targeted genome-editing delivery and a validated LDLR-null hepatocyte model for studying LDLR-dependent biology and disease.

## Introduction

Ionizable lipid nanoparticles (LNPs) have become the leading non-viral platform for therapeutic nucleic acid delivery, enabling approved mRNA vaccines, RNA interference medicines, and clinical programs for in vivo genome editing [1–3]. LNPs are multicomponent nanomaterials composed of an ionizable lipid, helper phospholipid, cholesterol, and PEG-lipid, which assemble around RNA payloads during controlled mixing. Their activity depends on a tightly balanced physicochemical profile. Ionizable lipids remain largely neutral at physiological pH, but become protonated in acidifying endosomes, promoting membrane destabilization and cytosolic RNA release [1,4,5]. Consequently, particle diameter, polydispersity, surface chemistry, apparent pKa, encapsulation efficiency, and RNA integrity must be controlled, because modest formulation changes can alter bio-distribution, endosomal escape, and editing potency [6,7].

The liver tropism of many conventional LNPs has been central to their success, but it also creates a biological limitation. After systemic exposure, LNPs adsorb plasma proteins and can acquire an apolipoprotein E (ApoE)-enriched corona that promotes hepatocyte uptake through LDLR and related lipoprotein receptors [8–10]. This endogenous pathway is efficient, yet it is not intrinsically cell selective within the liver and may be compromised when LDLR expression or function is reduced. Ligand-directed LNP engineering therefore offers a route to decouple delivery from passive protein-corona effects and redirect uptake through defined cell-surface interactions [9,11,12].

Aptamers are attractive targeting ligands for this purpose. These short single-stranded oligonucleotides are selected by SELEX to fold into target-binding structures with high affinity and specificity. Compared with antibodies, aptamers are smaller, synthetically accessible, chemically modifiable, and amenable to site-specific conjugation, making them well suited for nanoparticle surface functionalization [13–15]. Aptamer-decorated nanocarriers can engage cognate receptors and enter cells through receptor-mediated endocytosis, supporting selective delivery of drugs and nucleic acids to tumor cells [16,17]. TLS11a is a DNA aptamer selected against hepatocellular carcinoma cells and has been reported to bind HepG2 cells with high specificity, making it a rational ligand for directing LNPs to hepatocyte-derived cells through an LDLR-independent route [18,19].

This strategy is relevant to LDLR-driven disorders such as familial hypercholesterolemia (FH), a common inherited cause of premature atherosclerotic cardiovascular disease. In hepatocytes, LDLR mediates LDL binding and internalization, thereby regulating plasma LDL-cholesterol [20–25]. LDLR expression is controlled by cholesterol-sensitive feedback through the SCAP-SREBP-2 axis. When intracellular cholesterol falls, SCAP escorts SREBP-2 for proteolytic activation, inducing LDLR together with cholesterol biosynthesis and regulatory genes such as HMGCR, MVK, SQLE, and PCSK9 [26–30]. The clinical progress of LNP-delivered PCSK9 base-editing programs underscores the therapeutic potential of hepatic RNA-LNP genome editing and highlights the need for robust LDLR-deficient cellular models to evaluate delivery independently of LDLR function [31–35].

Here, we report the design and validation of a TLS11a-functionalized Cas9 mRNA/gRNA LNP platform and a matched LDLR-knockout HepG2 model. LNPs co-encapsulating Cas9 mRNA and an LDLR-targeting guide RNA were modified with a maleimide-terminated lipid anchor and conjugated to thiolated TLS11a. Orthogonal characterization confirmed nanoscale uniformity, preserved RNA cargo, retained ionizable pH-responsive behavior, and surface presentation of folded aptamer. Functionally, TLS11a decoration enhanced HepG2 mRNA delivery while reducing uptake in receptor-low control cells. In parallel, CRISPR disruption of LDLR exon 2 generated an LDLR-deficient HepG2 line validated at genomic, transcript, protein, and functional levels. This integrated system provides a modular platform for LDLR-independent targeted delivery and a hepatocyte-derived model for studying LDLR biology and therapeutic genome editing.

## Methods

### Cell lines and culture

Human hepatocellular carcinoma HepG2 cells and receptor-low Raji Burkitt lymphoma cells were cultured under standard conditions.. HepG2 cells (ATCC HB-8065) and Raji cells (ATCC CCL-86) were obtained from authenticated repository stocks, verified by short-tandem-repeat (STR) profiling, and confirmed mycoplasma-negative before use. HepG2 cells were maintained in Dulbecco’s modified Eagle’s medium, while Raji cells were maintained in RPMI-1640. Both media were supplemented with 10% heat-inactivated fetal bovine serum and 1% penicillin-streptomycin. Cells were incubated at 37 °C in a humidified atmosphere containing 5% CO_₂_. HepG2 cells were passaged at 70 to 80% confluence, and Raji cells were maintained in logarithmic growth.Cells were used between passages 3 and 20 (HepG2) and passages 3 and 25 (Raji). Raji B-lymphoblastoid cells, which lack hepatocyte-lineage markers and express low levels of the LDLR/ApoE hepatic-uptake axis, were included as a non-hepatic, receptor-low comparator to test cell-type-preferential delivery. Reagent suppliers and catalogue numbers are listed in Supplementary Table 1. All experiments were performed using mycoplasma-negative cells during exponential growth.

### Preparation of Cas9 mRNA/gRNA lipid nanoparticles

Cas9 mRNA/gRNA lipid nanoparticles were prepared by microfluidic mixing using established mRNA-LNP methods [1,2,6]. The lipid phase in ethanol contained the ionizable lipid, DSPC, cholesterol, and DMG-PEG2000 at a molar ratio of 50:10:38.5:1.5 [6,7]. For aptamer-conjugatable formulations, part of the PEG-lipid fraction was replaced with DSPE-PEG2000-maleimide to enable thiol-maleimide coupling [11,12]. The aqueous phase contained Cas9 mRNA and LDLR-targeting sgRNA in citrate buffer at pH 4.0. Lipid and aqueous streams were mixed at a 3:1 aqueous-to-ethanol flow-rate ratio, using an ionizable lipid-to-nucleic acid weight ratio of approximately 10:1 1 (corresponding to an amine-to-phosphate (N/P) ratio of approximately 6). The ionizable lipid used was SM-102 (Cayman Chemical), and Cas9 mRNA and LDLR-targeting sgRNA were co-encapsulated at a 4:1 mass ratio. Lipid and aqueous streams were combined in a microfluidic staggered-herringbone micromixer (NanoAssemblr, Precision NanoSystems) at a total flow rate of 12 mL/min at room temperature [1,6]. [1,6]. LNPs were diluted in PBS, purified by dialysis against PBS at pH 7.4, concentrated by ultrafiltration, passed through a 0.22 µm sterile filter, and stored at 4 °C. Dialysis was performed against PBS (pH 7.4) using a 100 kDa molecular-weight cut-off membrane. Purified formulations were protected from light and stored at 4 °C for up to 4 weeks before use.

### TLS11a Aptamer Functionalization of Cas9 mRNA/gRNA LNPs

The hepatocellular carcinoma-targeting DNA aptamer TLS11a [18] was synthesized with a 5′ thiol group and a spacer to support oriented surface conjugation. Before coupling, TLS11a was reduced with tris(2-carboxyethyl)phosphine to disrupt disulfide-linked aptamer dimers and generate free thiols. Reduction was performed with 10 mM TCEP at approximately 50-fold molar excess for 1 h at room temperature, and excess TCEP was removed by a desalting spin column (7 kDa MWCO) before refolding. The reduced aptamer was desalted and refolded in PBS containing 5 mM MgCl_₂_ by heating to 90 to 95 °C, followed by gradual cooling, as previously described [18,37]. Aptamer functionalization was performed on preformed maleimide-bearing LNPs through thiol-maleimide chemistry, producing a stable thioether linkage [12,37]. Residual maleimide groups were quenched, and unconjugated aptamer was removed by ultrafiltrationThiol-maleimide coupling was carried out at a 2:1 maleimide-to-thiol ratio for 2 h at room temperature followed by overnight incubation at 4 °C. Successful conjugation and removal of free aptamer were confirmed by the molecular-beacon hybridization assay and by XPS surface-elemental analysis described below. Non-decorated controls underwent the same handling steps without TLS11a. Throughout the study, bare LNPs refer to empty particles lacking Cas9 mRNA/gRNA and maleimide-PEG, non-decorated LNPs refer to cargo-loaded maleimide-bearing particles without aptamer, and TLS11a-LNPs refer to cargo-loaded particles after aptamer conjugation. This nomenclature separates effects of lipid composition, nucleic acid loading, maleimide incorporation, and aptamer-mediated targeting.

### Physicochemical characterisation

#### (i) Hydrodynamic Size and pH-Dependent Surface Charge Analysis

Hydrodynamic diameter and polydispersity index were measured by dynamic light scattering, while zeta potential was determined by laser Doppler electrophoresis in PBS at pH 7.4 and 25 °C. To assess ionizable charge-switching behavior, zeta potential was recorded across pH 4.0 to 7.4, and the apparent pKa was defined as the pH corresponding to charge neutralization [4,6]. All analyses were performed using at least three independently prepared LNP batches, with data reported as mean ± standard deviation.

#### (ii) Cryo-TEM Imaging of LNP Morphology

LNP formulations were deposited onto glow-discharged holey-carbon grids, blotted to form a thin aqueous film, and vitrified in liquid ethane using an automated plunge freezer. Samples were imaged under low-dose conditions on a cryogenic transmission electron microscope equipped with a direct electron detector [6].

#### (iii) mRNA Encapsulation and Serum Stability Assessment

mRNA encapsulation efficiency was measured using a RiboGreen assay, reported as total encapsulated nucleic acid (Cas9 mRNA plus sgRNA), by comparing fluorescence from intact and detergent-lysed LNPs (lysed with 2% (v/v) Triton X-100) [1,6]. Serum stability was assessed by incubating formulations in 50% serum (heat-inactivated fetal bovine serum) at 37 °C, followed by quantification of protected mRNA over 24 h at 0, 2, 6, and 24 h. Data were generated from independently prepared batches and reported as mean ± standard deviation.

### UV-Visible and Circular Dichroism Analysis of Nucleic Acid Loading and Structure

Cargo incorporation and nucleic acid folding were evaluated by scattering-corrected UV-visible spectroscopy and circular dichroism. Particle-derived light scattering was fitted and subtracted to resolve absorbance at 260 nm. UV-visible spectra were recorded from 220 to 350 nm against a matched buffer blank, and the scattering background was modelled as a power-law (Rayleigh-type) function fitted to the 320-350 nm region and subtracted to resolve the 260 nm nucleic acid peak [6]. Circular dichroism spectra were acquired across the far- and near-UV regions to assess helical and base-stacked nucleic acid structure. Thermal unfolding was monitored by variable-temperature circular dichroism from 25 to 95 °C. Circular dichroism spectra were acquired in PBS using a 1 mm path-length quartz cuvette at a scan speed of 50 nm/min, with mRNA, sgRNA, and aptamer at matched nucleic acid concentrations of approximately 5 µg/mL; thermal unfolding was monitored at 260 nm at a heating rate of 1 °C/min [38].

### Surface Elemental Analysis by X-ray Photoelectron Spectroscopy

Surface elemental composition was analyzed by X-ray photoelectron spectroscopy under ultra-high vacuum using a monochromatic X-ray source (Al Kα, 1486.6 eV). Spectra were charge-referenced to the adventitious C 1s peak at 284.8 eV to adventitious C 1s. Because XPS probes the outermost 5 to 10 nm of nanoparticles, nitrogen and phosphorus enrichment was used as evidence of peripheral aptamer presentation [39].

### Molecular Beacon Assay for TLS11a Surface Display

TLS11a surface presentation was confirmed using a fluorophore- and quencher-labeled molecular beacon complementary to the aptamer sequence. The beacon sequence is listed in Supplementary Table 1, and bare LNPs and non-decorated cargo-loaded LNPs were assayed separately as negative controls. The beacon was incubated with TLS11a-LNPs or control particles and hybridization to surface-accessible TLS11a relieved self-quenching, producing a concentration-dependent fluorescence signal (FAM excitation 495 nm, emission 520 nm; recorded on a fluorescence plate reader at 25 °C. Free TLS11a served as a reference to define the assay’s linear response range [40].

### Assessment of Aptamer-Mediated Cellular Uptake and Selectivity

Aptamer-mediated delivery was assessed using Cy5-labeled mRNA. HepG2 and Raji cells were incubated with bare, non-decorated, or TLS11a-functionalized LNPs under matched dosing and exposure conditions (Cy5-mRNA dose of 0.5 µg per well, corresponding to a total lipid concentration of approximately 10 µg/mL, for 6 h at 37 °C). The proportion of Cy5-positive cells was quantified by flow cytometry using at least 10,000 events per sample. Cellular internalization was visualized by confocal microscopy after DAPI nuclear counterstaining, with identical acquisition settings applied across all experimental groups.

### Guide RNA Design and Candidate Off-Target Evaluation

A single guide RNA targeting exon 2 of human LDLR, within the ligand-binding domain, was designed using established CRISPR design platforms (Benchling and CRISPOR, GRCh38/hg38 human genome build; the sgRNA spacer sequence is provided in **Supplementary Table 1**). Protospacers adjacent to a canonical PAM were ranked according to predicted on-target efficiency, sequence specificity, and genomic context [41,42]. Candidate off-target loci were computationally identified using 3- and 4-mismatch thresholds, then prioritized by risk score, with both DNA and RNA bulges included in the off-target search, coding or regulatory location, and proximity to biologically relevant genes. Experimentally detectable editing at selected sites was assessed by PCR amplification, Sanger sequencing and indel analysis. Editing at the intended LDLR locus was analyzed in parallel and indel spectra were deconvoluted from Sanger traces using ICE analysis (Synthego) to contextualize off-target activity relative to on-target disruption. Candidate off-target sites with measurable signal were flagged for confirmatory amplicon deep sequencing.

### CRISPR/Cas9 editing and isolation of LDLR-knockout clones

LDLR knockout was generated in HepG2 cells by delivery of Cas9 mRNA together with an sgRNA targeting exon 2 of LDLR (please include sgRNA sequence, here or in supplementary table) [41]. A fraction of the bulk-edited population was retained to quantify editing efficiency, while the remaining cells were plated by limiting dilution to derive single-cell clones. Expanded clones were screened by PCR amplification of the target locus followed by Sanger sequencing. Sequence traces were analyzed to define indel patterns, and alleles were classified as frameshift or in-frame relative to the protospacer. Clones carrying biallelic function-disrupting LDLR indels were selected for downstream molecular, protein, and functional validation.

### Quantification of Bulk Editing by T7 Endonuclease I Assay

Bulk LDLR editing efficiency was estimated using the T7 endonuclease I mismatch-cleavage assay [43,44]. The target locus was PCR (primer sequences, amplicon size, and cycling conditions are provided in **Supplementary Table 1**) from edited and untreated genomic DNA using primers designed to generate clearly resolvable cleavage products. Purified amplicons were denatured and gradually re-annealed to permit heteroduplex formation, then digestedwith 5 U T7 endonuclease I for 30 min at 37 °C with T7 endonuclease I. Cleavage products were separated by gel electrophoresison a 2% agarose gel and imaged under UV transillumination and editing efficiency was calculated from densitometric band intensities using the standard correction formula [43,44].

### RNA extraction and quantitative real-time PCR

Total RNA was isolated from experimental and control cells using a column-based RNA extraction kit (RNeasy, QIAGEN) and reverse-transcribed using a standardized cDNA synthesis protocol protocol (high-capacity cDNA reverse-transcription kit, Applied Biosystems). Quantitative real-time PCR was performed with gene-specific primers (listed in Supplementary Table 1) for *LDLR, HMGCR, MVK, SQLE, SREBF2, SREBF1, SCAP, and PCSK9*. Relative transcript abundance was normalized to a validated housekeeping gene *GAPDH* and calculated using the comparative Ct method across three independent biological replicates [45].

### Immunoblotting and immunofluorescence

Whole-cell lysates were separated by SDS-PAGE and transferred to membranes. Equal protein amounts (20 µg per lane) were resolved on 8-10% SDS-PAGE gels and transferred to PVDF membranes by wet transfer, then probed for LDLR, precursor SREBP-2, mature SREBP-2, and β-actin. Signals were detected using HRP-conjugated secondary antibodies and enhanced chemiluminescence. SREBP-2 processing was quantified by densitometry as the mature-to-precursor SREBP-2 ratio [26]. For immunofluorescence, fixed cells were stained for LDLR with a fluorophore-conjugated secondary antibody, counterstained with Hoechst, and imaged under identical acquisition settings. Cells were fixed in 4% paraformaldehyde and, for intracellular targets, permeabilized with 0.1% Triton X-100; surface LDLR staining was performed under non-permeabilized conditions. Densitometry was performed in ImageJ (Fiji), with target signals normalized to β-actin for all experimental groups to minimize acquisition bias. Antibody sources, catalog numbers, and dilutions are provided in the Supplementary Information.

### Lipoprotein Binding and Receptor-Mediated Uptake Assays

Receptor function was assessed using fluorescently labeled LDL, VLDL, and transferrin. To distinguish surface binding from internalization, cells were incubated with labeled LDL at 4 °C to measure binding or at 37 °C to permit uptake. Cells were serum-starved for 4 h before assay and incubated with fluorescently labelled LDL, VLDL, or transferrin (each at 10 µg/mL) for 1 h; after 4 °C binding, surface-bound ligand was distinguished from internalized signal by an acid/heparin wash [46]. After washing, fluorescence signal was quantified by microscopy and flow cytometry. VLDL uptake was measured in parallel. Transferrin uptake, mediated through the LDLR-independent transferrin receptor pathway, was included as a specificity control for preserved clathrin-mediated endocytosis [46]. Relative surface LDL-binding capacity was calculated from the 4 °C binding data.

### Cellular Cholesterol Quantification and Filipin Imaging

Total cellular cholesterol was measured using an enzymatic fluorometric assay and normalized to total protein content (measured by BCA protein assay). Free cholesterol was visualized by filipin III staining of fixed cells and quantified by fluorescence microscopy, with integrated signal intensity reported as fold change relative to control cells. Filipin III staining was performed for 30 min in the dark, and images were captured under matched UV excitation and exposure settings across groups [47].

### Statistical analysis

Data from at least three independent biological replicates are reported as mean ± standard deviation unless otherwise stated. Comparisons between two groups were performed using a two-tailed Student’s t-test, while multiple-group comparisons used one-way or two-way ANOVA with appropriate post hoc correction. Data normality was assessed using the Shapiro-Wilk test before parametric testing, and all t-tests were two-tailed. One-way ANOVA was used for single-factor comparisons and two-way ANOVA for experiments with two factors; post hoc correction was applied using Tukey’s test for all-pairwise comparisons, Dunnett’s test for comparisons against a single control, and Sidak’s test for pre-selected comparisons, as specified in each figure legend. The serum-stability time course was analysed by two-way repeated-measures ANOVA. For every panel, the exact n, the statistical test used, and the definition of the error bars (mean ± s.d.) are reported in the corresponding figure legend Exact p values are reported where applicable. Statistical significance was defined as p < 0.05, with significance annotations specified in each figure legend (\**p* < 0.05, \*\**p* < 0.01, \*\*\**p* < 0.001, \*\*\*\**p* < 0.0001; ns:, not significant).

## Results

### Physicochemical Characterization and Orthogonal Validation of TLS11a-Decorated Cas9 mRNA and gRNA LNPs

Cryo-electron microscopy demonstrated that unloaded LNPs were spherical, well dispersed, and relatively uniform, with an electron-dense core consistent with ionizable lipid and nucleic acid nanoparticle architecture (**Fig. 1a**). Co-encapsulation of Cas9 mRNA and LDLR-targeting gRNA preserved the overall morphology, while increasing internal electron density and particle diameter (**Fig. 1b**). Subsequent TLS11a aptamer functionalization produced intact, membrane-bounded particles with a modest increase in size and heterogeneity, a pattern consistent with surface-associated aptamer decoration rather than particle disruption (**Fig. 1c and Supplementary Fig. S1**). Dynamic light scattering confirmed a stepwise increase in Z-average diameter from 77 ± 7 nm for unloaded LNPs to 92 ± 8 nm after cargo loading and 105 ± 11 nm after TLS11a decoration (p < 0.01 for both comparisons; **Fig. 1d).** Size distributions remained narrow and clearly resolved across formulations (**Supplementary Fig. S1**). Polydispersity increased slightly from 0.12 ± 0.02 to 0.15 ± 0.02 after cargo loading (p < 0.05) and to 0.17 ± 0.02 after aptamer decoration, although this latter change was not significant. Importantly, all formulations retained a polydispersity index below 0.20, supporting acceptable colloidal uniformity (**Fig. 1e).** At physiological pH, zeta potential shifted progressively toward more negative values, from -3 ± 1.5 mV for unloaded LNPs to -7 ± 1.5 mV after cargo loading and -12 ± 2 mV after TLS11a decoration (**p < 0.01; Fig. 1f**). This shift is consistent with increasing surface exposure of anionic nucleic acid components, particularly the phosphate backbone of TLS11a. Across the pH titration, all formulations retained pH-responsive charge behavior, transitioning from approximately +38 mV at pH 4.0 to near neutrality at an apparent pKa of 6.3 to 6.5(the apparent pKa was determined by fitting the zeta-potential-versus-pH data to a sigmoidal Boltzmann function and taking the pH at which the fitted zeta potential equalled zero), within the endosomal pH range. Aptamer decoration preserved this profile while maintaining a slightly more negative surface charge at pH 7.4 (**Fig. 1g**). Cargo encapsulation remained high, decreasing only marginally from 92% to 89% (RiboGreen values refer to total encapsulated nucleic acid, i.e. Cas9 mRNA plus sgRNA) after TLS11a decoration, without statistical significance (**Fig. 1h**). Both cargo-loaded and TLS11a-decorated formulations were serum-stable, retaining approximately 88% and 85% of encapsulated mRNA, respectively, after 24 h in 50% serum at 37 °C (**Fig. 1i**). Because changes in size and zeta potential provide indirect evidence of aptamer coupling, surface display was further confirmed using molecular-beacon hybridization and XPS-based elemental analysis. TLS11a-complementary beacons generated concentration-dependent fluorescence on decorated particles but not on control LNPs, while XPS demonstrated increased surface nitrogen and phosphorus after aptamer addition **(Fig. 2d and Supplementary Fig. S1**).

**Figure 1.**
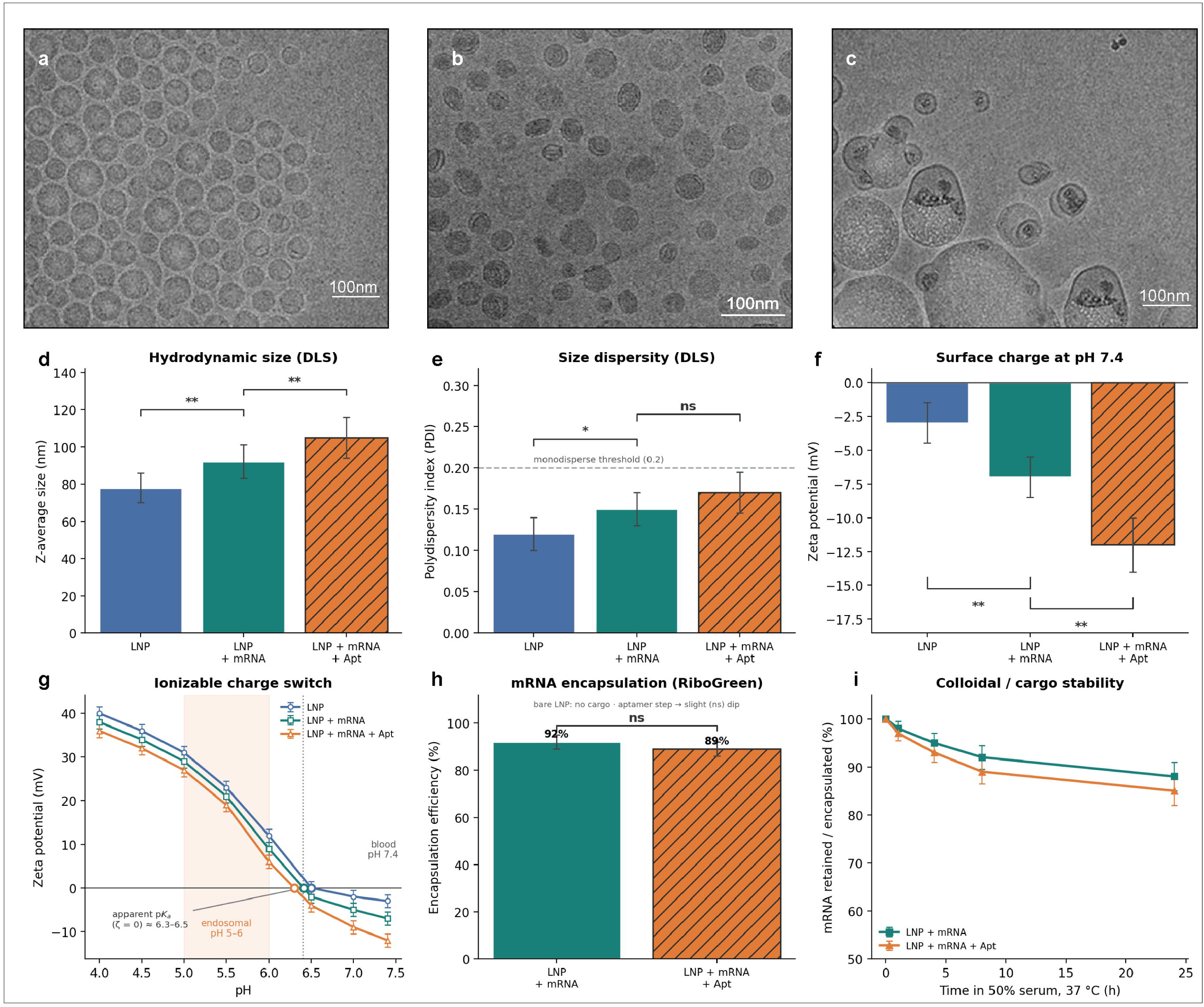
Physicochemical characterization of TLS11a-functionalized Cas9 mRNA/gRNA LNPs. Representative cryo-EM images of (a) bare LNPs, (b) Cas9 mRNA/gRNA-loaded LNPs, and (c) TLS11a-decorated cargo-loaded LNPs; scale bars, 100 nm. (d) Z-average diameter, (e) polydispersity index, and (f) zeta potential at pH 7.4. (g) pH-dependent zeta-potential titration showing ionizable charge switching, apparent pKa at ζ = 0, and the endosomal pH range. (h) mRNA encapsulation efficiency measured by RiboGreen. (i) Retention of encapsulated mRNA during incubation in 50% serum at 37 °C for 24 h. Data are mean ± s.d. from at least three independent batches; *p < 0.05, **p < 0.01; ns, not significant.

**Figure 2.**
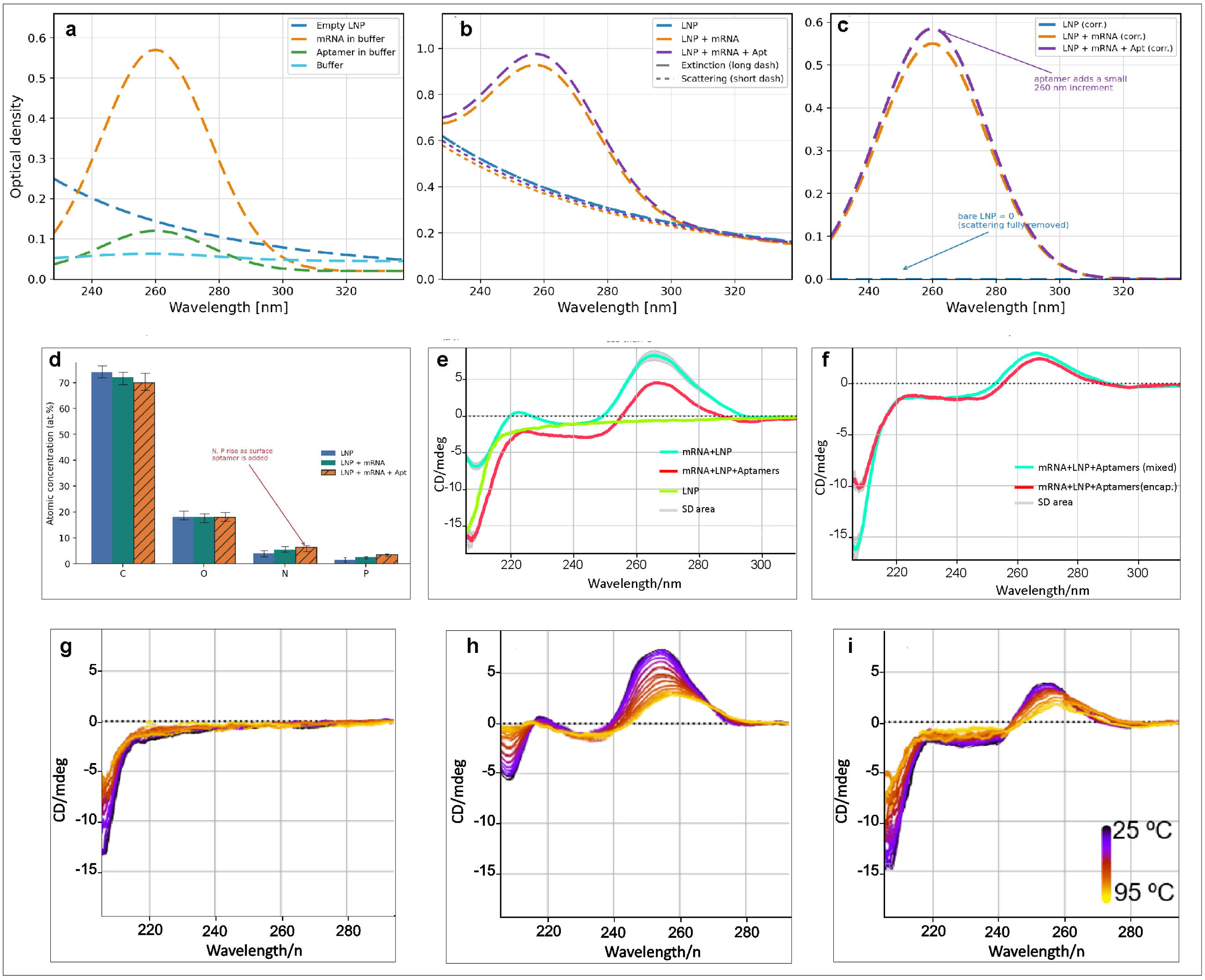
Spectroscopic and surface validation of TLS11a-functionalized Cas9 mRNA/gRNA LNPs. (a) UV absorbance spectra of individual components, including empty LNPs, mRNA in buffer, free TLS11a aptamer, and buffer alone, confirming that the 260 nm signal is nucleic acid derived. (b) Raw extinction and fitted scattering spectra of bare LNPs, Cas9 mRNA/gRNA-loaded LNPs, and TLS11a-functionalized cargo-loaded LNPs, showing a nucleic acid absorbance peak superimposed on the LNP scattering background in cargo-containing formulations. (c) Scattering-corrected UV spectra showing removal of the bare-LNP background and a modest 260 nm increase after TLS11a functionalization. (d) XPS surface atomic composition, with an approximate sampling depth of 5 to 10 nm, showing increased surface nitrogen and phosphorus after aptamer addition, consistent with peripheral oligonucleotide enrichment. (e) CD spectra showing ordered nucleic acid structure in cargo-loaded formulations, with a positive band near 260 nm and a negative far-UV band. Functional accessibility of TLS11a was assessed separately by molecular-beacon hybridization in Supplementary Fig. S1. (f) CD comparison of mixed versus encapsulated cargo in TLS11a-functionalized LNPs, supporting preservation of the dominant secondary-structure signature after encapsulation. (g to i) Variable-temperature CD spectra from 25 to 95 °C for (g) bare LNPs, (h) TLS11a-functionalized cargo-loaded LNPs, and (i) cargo-loaded LNPs. Cargo-containing formulations showed progressive loss of the 260 nm band during heating. Shaded regions indicate s.d. from at least three independent formulation batches.

### Orthogonal Spectroscopic Validation of Nucleic Acid Incorporation, Surface Aptamer Display, and Structural Retention

To verify nucleic acid incorporation and structural preservation, bare LNPs, Cas9 mRNA/gRNA-loaded LNPs, and TLS11a-functionalized cargo-loaded LNPs were analyzed by scattering-corrected UV-visible spectroscopy and circular dichroism. Component spectra confirmed that the dominant 260 nm absorbance originated from nucleic acid cargo, with free mRNA producing a strong signal, free TLS11a generating a smaller oligonucleotide peak, and empty LNPs contributing mainly to the rising ultraviolet scattering background (**Fig. 2a**). In intact formulations, raw extinction spectra showed a clear 260 nm nucleic acid peak superimposed on the LNP scattering profile in both cargo-loaded groups, whereas bare LNPs lacked a discrete nucleic acid absorbance peak (**Fig. 2b**). After mathematical subtraction of the scattering component, the bare-LNP spectrum returned close to baseline, while cargo-loaded formulations retained a defined 260 nm peak. The TLS11a-functionalized formulation showed a modest additional 260 nm increase, consistent with contribution from the surface-conjugated aptamer (**Fig. 2c**). Surface-sensitive XPS provided independent support for aptamer display at the nanoparticle exterior. Because XPS probes only the outer 5 to 10 nm, the observed increase in surface nitrogen and phosphorus after TLS11a addition was consistent with enrichment of a phosphate-backbone oligonucleotide at the particle periphery (**Fig. 2d).** This interpretation should be strengthened by reporting N/P ratios or fold enrichment relative to non-functionalized LNPs.

Circular dichroism further supported retention of ordered nucleic acid structure within the formulations. Bare LNPs produced minimal signal in the nucleic acid region, whereas cargo-loaded formulations displayed a positive band near 260 nm and a negative band in the far-UV region, consistent with base stacking and helical secondary structure **(Fig. 2e).** The stronger signal in TLS11a-functionalized LNPs likely reflects the additional contribution of surface-displayed aptamer. Because bulk CD is dominated by encapsulated mRNA/gRNA and cannot alone confirm functional TLS11a folding, target-accessible aptamer presentation was independently supported by molecular-beacon hybridization (**Supplementary Fig. S1**). To assess whether encapsulation altered nucleic acid conformation, CD spectra were compared between formulations containing mixed versus encapsulated mRNA/gRNA cargo. The near-overlapping spectra indicate that the formulation process preserved the dominant secondary-structure signature while enabling the serum protection shown in Fig. 1i **(Fig. 2f**). Variable-temperature CD from 25 to 95 °C showed no structured thermal transition for bare LNPs, whereas cargo-containing formulations exhibited progressive loss of the 260 nm band during heating **Fig. 2h and 2i** supporting cooperative unfolding of ordered nucleic acid structure rather than nonspecific random-coil behavior.

### TLS11a Functionalization Enhances Preferential mRNA Delivery to HepG2 Cells

After confirming TLS11a surface display and cargo integrity, we assessed whether aptamer functionalization improved cell-selective mRNA delivery. Cy5-labeled mRNA uptake was quantified by flow cytometry and visualized by confocal microscopy using HepG2 cells and Raji cells as a non-hepatic comparator (**Supplementary Fig. S2**). Non-decorated cargo-loaded LNPs achieved detectable delivery in HepG2 cells, with 39% Cy5-positive cells. Values are the mean of three independent experiments; group comparisons used a two-tailed Student’s t-test, with exact p values and error bars defined in the figure legend. TLS11a functionalization increased the Cy5-positive HepG2 fraction to 79%, indicating an approximate two-fold enhancement over non-decorated LNPs under matched exposure conditions. The same TLS11a-functionalized formulation produced 31% Cy5-positive Raji cells, supporting preferential delivery to HepG2 cells rather than a uniform increase in cellular uptake. The LNP-only condition showed minimal background fluorescence, (here, “LNP-only” denotes empty, cargo-free LNPs used as a fluorescence-background control), confirming low nonspecific Cy5 signal in the assay. Confocal microscopy supported the flow-cytometry findings, showing the strongest Cy5-associated signal in TLS11a-functionalized HepG2 cells, weaker signal in non-decorated HepG2 and TLS11a-treated Raji cells, and minimal signal in the LNP-only control. DAPI staining confirmed the presence of comparable cell fields across conditions. Collectively, these data support TLS11a-associated enhancement of mRNA delivery to HepG2 cells. However, because scrambled-aptamer and free-aptamer competition controls were not included, these results should be interpreted as preferential, aptamer-associated delivery rather than definitive sequence-specific targeting.

### CRISPR/Cas9 Disruption of LDLR Exon 2 Establishes a Multilevel-Validated LDLR-Knockout HepG2 Model

To establish an LDLR-deficient HepG2 model, we designed a single guide RNA targeting exon 2 of LDLR within the ligand-binding domain, where frameshift-inducing indels were expected to disrupt receptor function **(Fig. 3a to c).** The gRNA target site and PAM were mapped relative to the LDLR coding sequence, confirming disruption within an early functional region of the receptor **(Fig. 3d).** Sanger sequencing of single-cell-derived clones identified biallelic deletions in three validated clones: Clone 1, -10/-13 bp; Clone 2, -5/-11 bp; and Clone 3, -11/-7 bp. These deletions were predicted to introduce frameshifts immediately downstream of the protospacer, supporting functional disruption of LDLR. On-target genome editing was further assessed by T7 endonuclease I mismatch-cleavage analysis. Amplification of the target region yielded an expected 563 bp product, with cleavage products of 400 bp and 163 bp detected after mismatch digestion, consistent with indel formation at the LDLR target site **(Fig. 3e).** Bulk editing efficiency was estimated at 68% by densitometric quantification of T7E1 cleavage products (standard correction formula; n=3 independent experiments), with replicate-level quantification provided in Supplementary **Fig. S3**. Following single-cell cloning and expansion, three biallelic LDLR-disrupted clones were recovered and prioritized for molecular and cellular validation. Loss of LDLR expression was confirmed at transcript, protein, and cellular levels. LDLR mRNA was significantly reduced in edited cells compared with control cells (n=3 independent biological replicates; two-tailed Student’s t-test) **(p = 1.3 × 10**^-^**²; Fig. 3f)**, consistent with disruption of the coding sequence and possible nonsense-mediated decay. Immunoblotting demonstrated loss of detectable LDLR protein in the knockout population, while β-actin confirmed comparable protein loading (Fig. 3g). Immunofluorescence further supported receptor ablation at the single-cell level. Control HepG2 cells showed strong LDLR staining, whereas LDLR-knockout cells showed markedly reduced signal under matched imaging conditions. Hoechst staining confirmed comparable nuclear fields across conditions, indicating that loss of LDLR signal was not attributable to reduced cell number (**Fig. 3h to l)**. Together, these data support successful generation of a genetically disrupted and molecularly validated LDLR-knockout HepG2 model.

**Figure 3.**
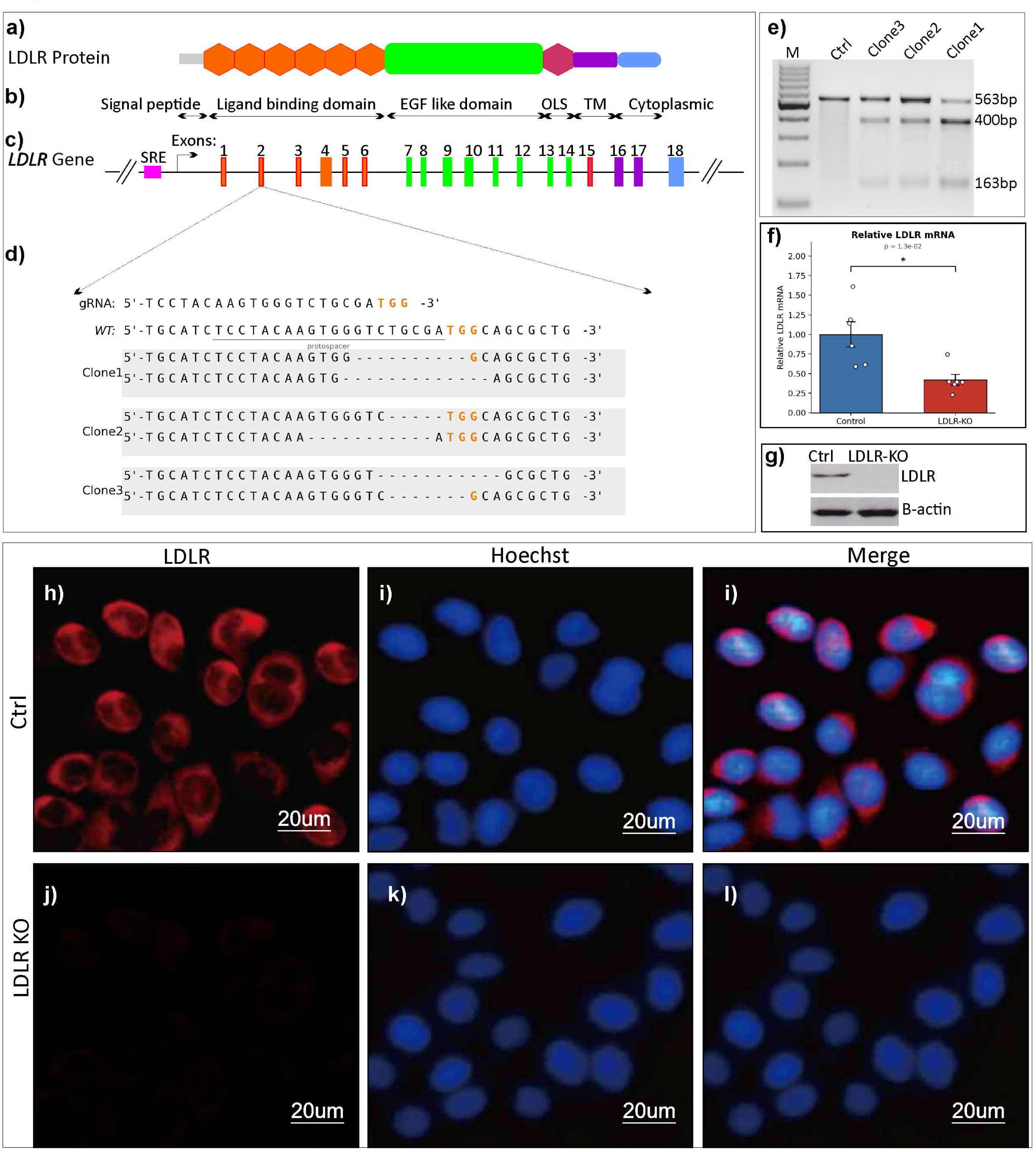
Genomic and molecular validation of LDLR-knockout HepG2 cells. (a) Schematic of LDLR protein domains. (b, c) LDLR gene organization, including the sterol-regulatory element and exons 1 to 18, showing sgRNA targeting of exon 2 within the ligand-binding domain. (d) Sanger-confirmed sequences of the wild-type locus and three edited clones, with the protospacer and PAM indicated and deletion sizes shown. Clones 2 and 3 contain frameshift alleles, whereas Clone 1 contains in-frame deletions within the ligand-binding domain. Full sequence traces are provided in Supplementary Fig. S3. (e) T7E1 mismatch-cleavage assay showing the 563 bp amplicon and expected 400 bp and 163 bp cleavage products; editing quantification is provided in Supplementary Fig. S3. (f) Relative LDLR mRNA expression in control and LDLR-KO cells. (g) LDLR immunoblot with β-actin loading control. (h to l) LDLR immunofluorescence, Hoechst nuclear staining, and merged images in control and LDLR-KO HepG2 cells; scale bars, 20 µm.

### LDLR knockout markedly impairs LDL binding and uptake

To define the functional impact of LDLR disruption on lipoprotein handling, fluorescent LDL binding and uptake were compared between control and LDLR-KO HepG2 cells under temperature-controlled conditions. In control cells, labeled LDL produced strong surface-associated fluorescence at 4 °C, where internalization is minimized, and clear intracellular punctate signal at 37 °C, consistent with receptor binding followed by endocytic uptake **(Fig. 4a)**. In contrast, LDLR-KO cells showed markedly weaker LDL signal at both temperatures, indicating impaired surface binding and reduced internalization rather than complete elimination of all residual lipoprotein association **(Fig. 4b)**. Quantitative analysis confirmed a major reduction in LDL uptake, with LDLR-KO cells retaining only approximately 15% of the control signal **(p = 6.8 × 10**⁻**¹□ ; Fig. 4c)**. Relative LDL binding capacity measured at 4 °C was also significantly reduced **(p = 1.9 × 10**⁻³; **Fig. 4f)**, supporting loss of functional surface LDLR activity.

**Figure 4.**
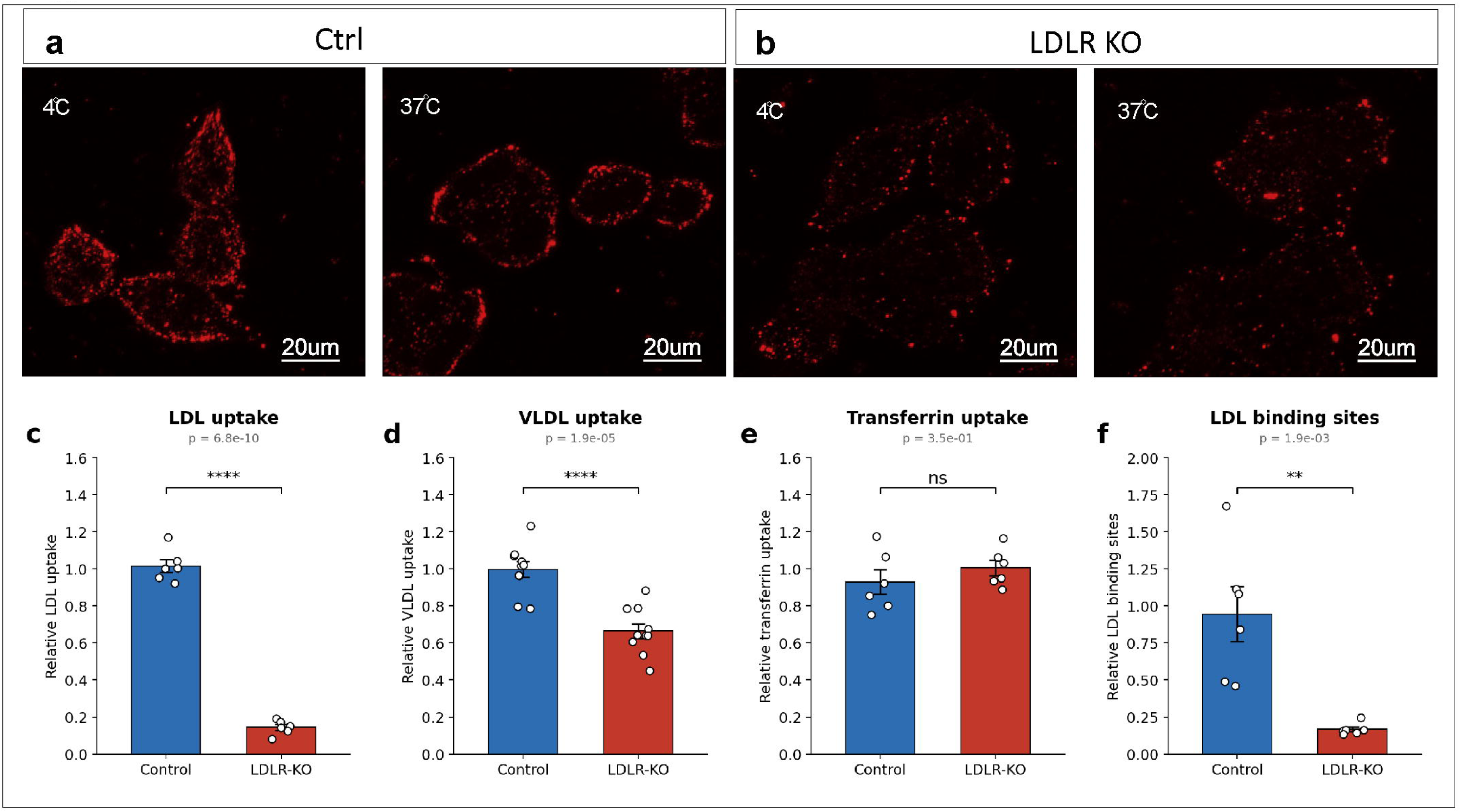
**Functional LDLR deficiency demonstrated by lipoprotein binding and uptake assays.**(a, b) Fluorescence microscopy of labeled LDL surface binding at 4 °C and internalization at 37 °C in (a) control and (b) LDLR-KO HepG2 cells; scale bars, 20 µm. (c) Relative LDL uptake, (d) relative VLDL uptake, (e) relative transferrin uptake, and (f) relative surface LDL-binding capacity in control and LDLR-KO cells. LDLR knockout markedly reduced LDL uptake and surface binding, partially reduced VLDL uptake, and preserved transferrin receptor-mediated uptake. Bar graphs show mean ± s.d. with individual replicates overlaid; exact p values are shown in each panel. ns: not significant.

To evaluate pathway specificity, uptake of VLDL and transferrin was measured in parallel. VLDL uptake was significantly reduced in LDLR-KO cells, although the decrease was more modest than for LDL, reaching approximately 33% below control levels **(p = 1.9 × 10**⁻□ ; **Fig. 4d)**. This pattern is consistent with the broader receptor repertoire involved in hepatic VLDL and remnant clearance, including LDLR-related protein 1, scavenger receptor class B type I, and heparan sulfate proteoglycans. By contrast, transferrin uptake, mediated through the transferrin receptor pathway, was not significantly altered in LDLR-KO cells **(p = 0.35; Fig. 4e)**. This preserved transferrin signal supports maintenance of transferrin receptor-dependent endocytosis, although it should not be interpreted as proof that all clathrin-mediated pathways are unchanged. The data therefore separate an LDLR-dependent defect in lipoprotein handling from nonspecific impairment of membrane trafficking or cell viability under these assay conditions. Together, the profound loss of LDL binding and uptake, partial reduction in VLDL uptake, and preserved transferrin uptake demonstrate that CRISPR-mediated LDLR disruption produces a functionally validated LDLR-deficient HepG2 model rather than a generalized defect in cellular endocytosis.

### LDLR Knockout Triggers Intracellular Cholesterol Depletion and Compensatory SCAP-SREBP-2 Activation

To determine whether loss of LDLR-mediated cholesterol import produces a compensatory sterol response, we measured cellular cholesterol content and activation of the SCAP-SREBP-2 pathway in control and LDLR-KO HepG2 cells. Consistent with the impaired LDL binding and uptake shown in Fig. 4, LDLR-KO cells contained significantly less total cellular cholesterol and showed reduced filipin signal, indicating depletion of both total and free cholesterol pools (total cholesterol was measured as the sum of free and esterified cholesterol) **(Fig. 5a,b)**. The parallel reduction in biochemical cholesterol content and filipin fluorescence supports a direct metabolic consequence of LDLR disruption in HepG2 cells across independent assay readouts. Cholesterol depletion was accompanied by coordinated activation of sterol-regulatory transcriptional programs. LDLR-KO cells showed increased expression of SREBF2 and its escort factor SCAP, whereas SREBF1 remained unchanged, supporting a response preferentially linked to cholesterol rather than fatty-acid regulation **(Fig. 5f to h)**. Downstream cholesterol biosynthesis genes, including HMGCR, MVK, and SQLE, were also significantly upregulated, consistent with a compensatory attempt to restore intracellular sterol availability **(Fig. 5c to e)**. This pattern is biologically coherent because reduced receptor-mediated cholesterol entry would be expected to increase endogenous cholesterol synthesis through SREBP-2-dependent transcriptional regulation.

**Figure 5.**
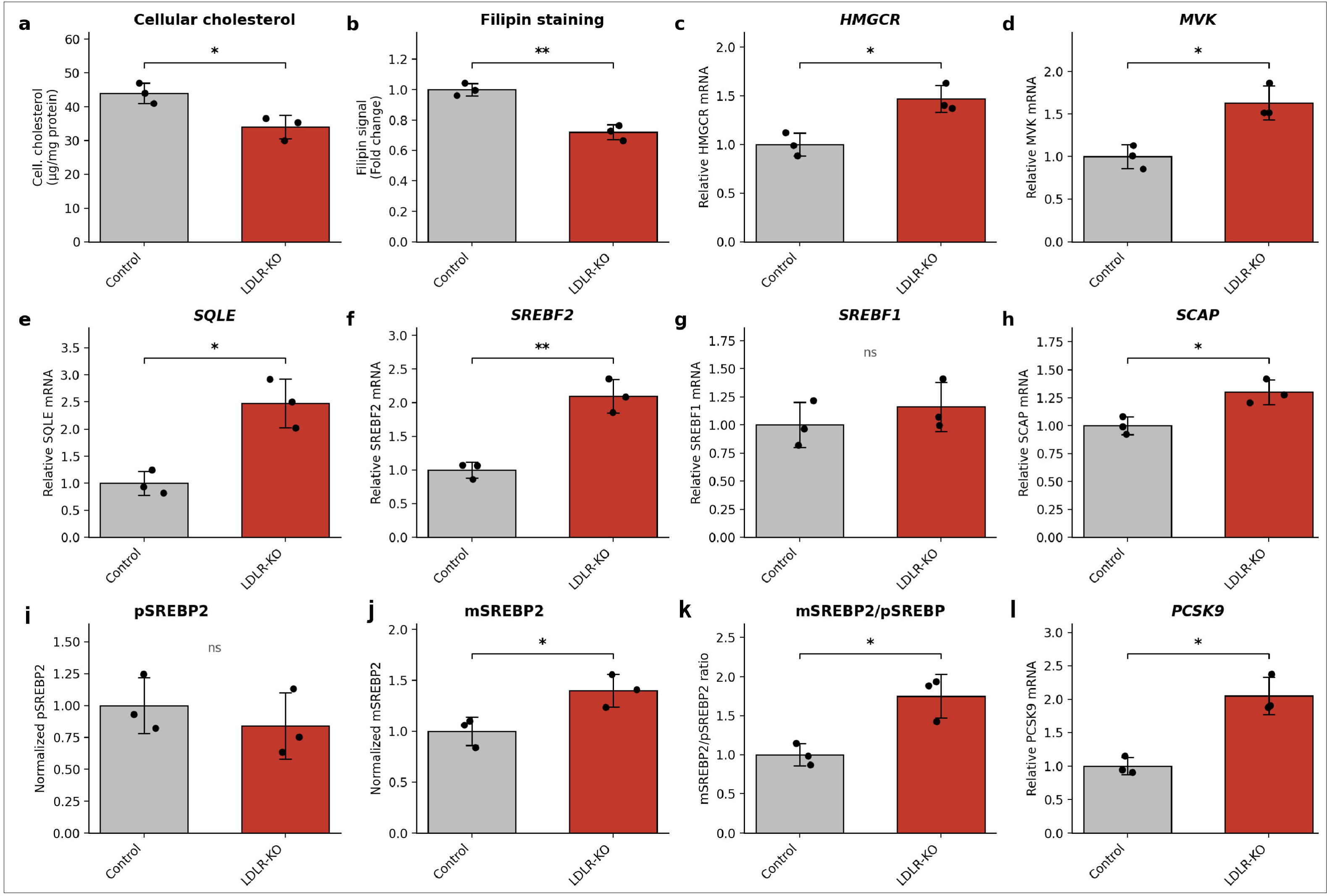
LDLR knockout depletes intracellular cholesterol and activates the SCAP–SREBP-2 feedback axis. (a) Total cellular cholesterol normalized to protein and (b) filipin-based free cholesterol signal in control and LDLR-KO HepG2 cells. (c to e) Relative mRNA expression of cholesterol biosynthesis genes HMGCR, MVK, and SQLE. (f to h) Relative expression of SREBF2, SREBF1, and SCAP, assessing sterol-regulatory specificity. (i to k) Quantification of SREBP-2 processing, including precursor pSREBP2, mature mSREBP2, and the mSREBP2/pSREBP2 ratio. (l) Relative PCSK9 mRNA, a canonical SREBP-2 target. Bars show mean ± s.d. with individual biological replicates overlaid, n = 3 per group. Panel-specific p values are shown, and statistical tests are described in Methods. Significance is indicated for all panels as ns, not significant; *p < 0.05; **p < 0.01.

Because SREBP-2 activity is regulated mainly through sterol-sensitive proteolytic processing, we next assessed precursor and mature SREBP-2 protein abundance. Precursor SREBP-2 was not significantly altered, whereas mature SREBP-2 increased in LDLR-KO cells, leading to an elevated mature-to- precursor SREBP-2 ratio (precursor and mature SREBP-2 band intensities were each normalized to β-actin before computing the mature-to-precursor ratio) **(Fig. 5i to k)**. This pattern supports enhanced SREBP-2 processing rather than a simple increase in total precursor abundance. In agreement with increased SREBP-2 activity, PCSK9 mRNA was also elevated in LDLR-KO cells **(Fig. 5l)**. However, because LDLR is genetically disrupted in this model, increased PCSK9 expression is unlikely to restore LDLR-dependent uptake and should be interpreted primarily as a marker of pathway activation. Together, these data show that LDLR knockout reduces intracellular cholesterol availability and robustly activates a coherent SCAP-SREBP-2 feedback response, characterized by increased cholesterol biosynthesis gene expression, enhanced SREBP-2 maturation, and induction of PCSK9. This integrated transcriptional, biochemical, and protein-processing profile provides functional evidence that CRISPR-mediated LDLR disruption produces a biologically meaningful cholesterol-import defect in HepG2 cells, while preserving the expected compensatory sterol-sensing machinery under these experimental conditions in this cellular model.

## Discussion

This study developed and evaluated a TLS11a-functionalized ionizable LNP platform intended to support cell-selective delivery of CRISPR/Cas9 cargo while reducing reliance on the endogenous ApoE-LDLR uptake route. In parallel, we generated an LDLR-deficient HepG2 model to provide a stringent cellular context in which LDLR-dependent lipoprotein handling, sterol feedback, and LDLR-independent delivery could be interrogated. The main finding is that TLS11a decoration can enhance preferential mRNA delivery to HepG2 cells without compromising core LNP physicochemical properties, while CRISPR-mediated LDLR disruption creates a functionally validated model of impaired LDL uptake and compensatory SCAP-SREBP-2 activation.

The physicochemical results support a coherent stepwise assembly process. Cryo-electron microscopy showed intact, membrane-bounded particles, and progression from bare to cargo-loaded to TLS11a-decorated LNPs was accompanied by a stepwise increase in Z-average diameter from 77 to 92 to 105 nm, while the polydispersity index remained below 0.20, consistent with acceptable colloidal uniformity [1,6]. This pattern is compatible with nucleic acid loading followed by addition of a hydrated surface aptamer layer, although morphology and size alone cannot prove surface presentation. Zeta potential shifted progressively toward more negative values at physiological pH, from -3 to -7 to -12 mV, consistent with increasing exposure of anionic nucleic acid components, particularly the phosphate backbone of TLS11a [9,12]. Importantly, the formulations retained a pH-responsive charge transition, with an apparent pKa of approximately 6.3 to 6.5, within the range associated with endosomal ionization and cytosolic delivery of ionizable LNP systems [7,49]. The minimal change in encapsulation efficiency after aptamer decoration, from 92% to 89%, together with retention of encapsulated mRNA in serum, indicates that surface functionalization was achieved without major destabilization of the cargo compartment [6].

Because size and charge changes provide only indirect evidence of conjugation, aptamer display was assessed using complementary analytical methods. Scattering-corrected UV spectroscopy resolved the expected 260 nm nucleic acid absorbance in cargo-loaded formulations, with a modest additional signal after TLS11a decoration that was consistent with contribution from the surface oligonucleotide [6]. Circular dichroism showed a positive band near 260 nm and a negative far-UV band, supporting preservation of ordered nucleic acid secondary structure within the formulations [38]. However, CD is dominated by the encapsulated mRNA/gRNA and cannot, by itself, confirm the functional tertiary fold of a surface aptamer. For this reason, molecular-beacon hybridization provided an important accessibility assay, showing a concentration-dependent signal on TLS11a-decorated particles but not on bare controls [40]. X-ray photoelectron spectroscopy further supported peripheral aptamer enrichment by showing increased surface nitrogen and phosphorus within the outer 5 to 10 nm of the particle surface [39]. Together, these optical, hybridization-based, and surface elemental analyses support, but should not overstate, successful TLS11a presentation on the LNP exterior.

The delivery data indicate that TLS11a functionalization enhances preferential uptake in HepG2 cells. TLS11a-decorated LNPs increased the Cy5-positive HepG2 fraction from 39% with non-decorated LNPs to 79%, while the same decorated formulation produced 31% Cy5-positive Raji cells. This pattern supports cell-type-preferential delivery rather than absolute HepG2 specificity. Confocal microscopy was concordant with the flow-cytometry data, showing the strongest cytoplasmic Cy5 signal in TLS11a-functionalized HepG2 cells and weaker signal in non-decorated HepG2 and TLS11a-treated Raji cells [16,17]. These findings are consistent with the broader principle that engineered surface ligands can redirect LNP uptake away from passive corona-driven behavior and toward ligand-associated cell recognition [8–10]. Nevertheless, the mechanistic interpretation should remain cautious. In this study, receptor-independent delivery means reduced dependence on the ApoE-LDLR pathway, not absence of receptor involvement. The precise TLS11a-binding target on HepG2 cells remains incompletely defined, and sequence-specific targeting should be further tested using scrambled aptamer controls, free TLS11a competition, and direct internalization assays. It is also important to distinguish the two experimental arms of the work: the LDLR-knockout HepG2 line was generated using conventional Cas9 mRNA/gRNA delivery, whereas the TLS11a-LNP system was evaluated using Cy5-mRNA uptake. Demonstrating efficient genome editing directly from TLS11a-functionalized Cas9 mRNA/gRNA LNPs will be essential before claiming a fully integrated targeted-editing platform.

The LDLR-editing arm provides an experimentally useful, genetically defined hepatocyte model. Targeting exon 2 was biologically appropriate because this region lies within the ligand-binding domain, where disruptive indels are expected to impair LDL interaction. T7 endonuclease I mismatch-cleavage analysis confirmed on-target editing in the edited population, yielding the expected 563 bp amplicon and 400 and 163 bp cleavage products, with an estimated bulk editing efficiency of 68% [43,44]. Because T7E1 is semi-quantitative and can under-detect certain indel patterns, sequencing-based validation remains critical [44]. Sanger sequencing of isolated clones confirmed target-site deletions, and the combined transcript, protein, immunofluorescence, and uptake data established functional loss of LDLR activity [31,44]. The distinction between bulk editing efficiency and recovered clone distribution is important, because clone recovery after single-cell expansion does not necessarily reflect the true composition of the original edited population. For translational rigor, amplicon deep sequencing and unbiased off-target analysis would further strengthen the genomic characterization.

Functional uptake experiments confirmed that LDLR disruption produces a selective lipoprotein-handling defect rather than a generalized endocytic failure. LDL uptake was reduced by approximately 85%, and surface LDL-binding capacity decreased by a similar magnitude, consistent with loss of the receptor responsible for high-affinity LDL binding [24,25,46]. Transferrin uptake was unchanged, indicating preserved transferrin receptor-mediated uptake under the tested conditions [46]. This result should be interpreted as preservation of this control pathway, rather than proof that every clathrin-mediated route remains intact. VLDL uptake was reduced more modestly, by approximately 33%, which is biologically plausible because VLDL and remnant clearance involve multiple hepatic receptors and matrix-associated pathways. The residual LDL signal and partial preservation of VLDL uptake are consistent with contributions from LDLR-independent routes, including heparan-sulfate proteoglycans, LRP1, and SR-BI [50–53]. These pathways were not individually blocked in the present study, so their involvement is inferred from established hepatic lipoprotein biology rather than directly dissected here.

The cholesterol-homeostasis data provide a second layer of functional validation. Loss of LDLR-mediated lipoprotein entry reduced total and free cellular cholesterol and activated the SCAP-SREBP-2 feedback pathway [26–28]. LDLR-deficient cells showed increased SREBF2 and SCAP expression, unchanged SREBF1, and induction of cholesterol-biosynthesis genes including HMGCR, MVK, and SQLE [27,30]. This pattern supports a sterol-focused response rather than a broad, nonspecific transcriptional stress program. Protein-level analysis further strengthened this interpretation: precursor SREBP-2 was not significantly altered, whereas mature SREBP-2 increased, raising the mSREBP2/pSREBP2 ratio and supporting enhanced sterol-responsive processing [26,28]. PCSK9, a canonical SREBP-2 target, was also induced [29]. In this LDLR-deficient background, however, increased PCSK9 expression is unlikely to restore LDLR-dependent uptake. More broadly, the model illustrates an intact sterol-sensing circuit whose principal import effector is genetically disabled, a configuration relevant to LDLR-deficient disease biology and to the limited utility of interventions that require functional LDLR expression [29,34].

Several limitations should guide interpretation and future work. First, all experiments were performed in cultured cells, and in vivo performance of TLS11a-decorated LNPs may be shaped by serum nuclease exposure, protein corona formation, immune recognition, biodistribution, ligand density, aptamer orientation, and endosomal escape [5,10,13,37]. Second, the delivery experiment measured Cy5-mRNA uptake rather than genome editing from the TLS11a-functionalized Cas9 mRNA/gRNA formulation. Third, LDLR-independent delivery and LDLR-independent lipoprotein uptake were not mechanistically resolved at the receptor level. Finally, unbiased off-target profiling and deeper clone sequencing would be required before translational claims can be made [42,44].

## Conclusion

In this study, we assembled a TLS11a-functionalized ionizable LNP platform for CRISPR/Cas9 cargo and, in parallel, generated a multilevel-validated LDLR-knockout HepG2 model. Rather than introducing new targeting ligands or delivery chemistries, the work integrates established components, a hepatocellular carcinoma–selective aptamer, thiol-maleimide conjugation, and microfluidic mRNA-LNP formulation into a single defined system characterized by a complete orthogonal workflow. Across cryo-electron microscopy, dynamic light scattering, pH-dependent zeta-potential titration, UV and circular dichroism spectroscopy, X-ray photoelectron spectroscopy, and molecular-beacon hybridization, TLS11a decoration preserved the features expected of a functional ionizable RNA-LNP: nanoscale size, colloidal uniformity, an apparent pKa within the endosomal-ionization range, high encapsulation, and serum stability. Consistent with prior aptamer-nanoparticle studies, functionalization enhanced Cy5-mRNA uptake in HepG2 cells relative to non-decorated particles and a non-hepatic comparator. Because sequence-specificity controls were not included, these data reflect ligand-associated, cell-preferential delivery rather than receptor-specific targeting. Separately, CRISPR disruption of LDLR exon 2 produced a genetically defined HepG2 line validated at genomic, transcript, protein, immunofluorescence, and functional levels. In agreement with established models of hepatic LDLR loss, it showed markedly reduced LDL binding and uptake, partially preserved VLDL handling, unaffected transferrin uptake, and a coherent compensatory SCAP-SREBP-2 response confirming a selective lipoprotein-handling defect rather than generalized endocytic failure.

The principal value of this work is therefore methodological and integrative; a reproducible, orthogonally characterized aptamer-LNP formulation and a rigorously validated LDLR-null hepatocyte model for studying LDLR-independent delivery and cholesterol-regulatory biology. Its central limitation is equally clear editing was demonstrated using conventional Cas9 mRNA/gRNA delivery, whereas the TLS11a-LNP was evaluated only by Cy5-mRNA uptake. Demonstrating efficient, sequence-specific editing directly from the functionalized formulation, with appropriate controls, off-target profiling, and in vivo evaluation, will be required before this system can advance as an integrated targeted genome-editing platform.

## Supplementary Figure Legends

**Supplementary Figure S1.**
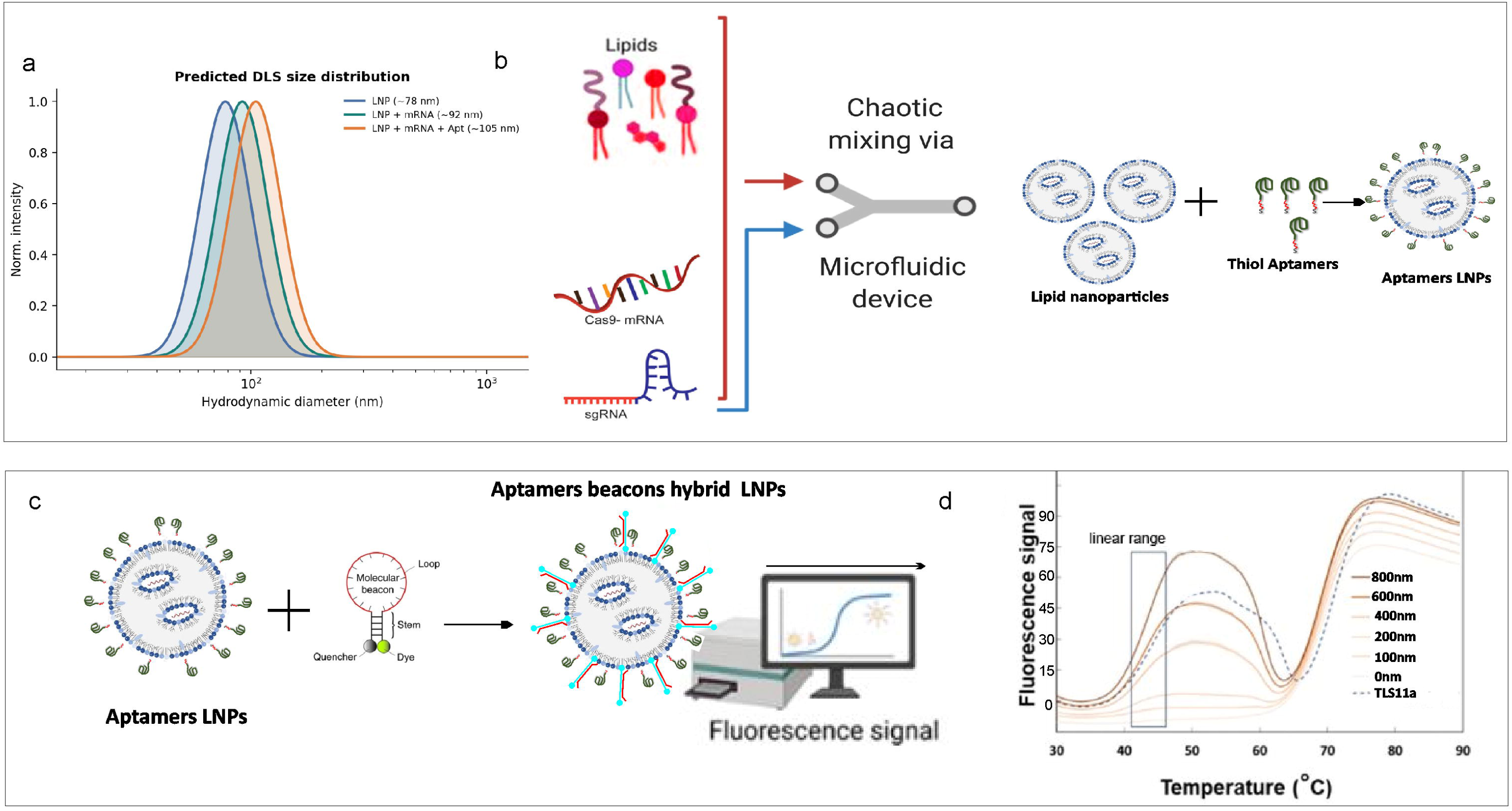
Microfluidic LNP assembly and molecular-beacon validation of TLS11a surface display. (a) DLS size-distribution overlay for bare LNPs, Cas9 mRNA/gRNA-loaded LNPs, and TLS11a-functionalized cargo-loaded LNPs, showing a stepwise increase in hydrodynamic diameter and narrow particle-size distributions that support preserved colloidal quality. (b) Schematic of microfluidic formulation, in which ionizable lipid components are rapidly mixed with Cas9 mRNA/gRNA to generate cargo-loaded LNPs, followed by conjugation with thiol-modified TLS11a aptamers to produce surface-functionalized particles. (c) Molecular-beacon assay principle showing hybridization of TLS11a-complementary beacons to surface-accessible TLS11a on decorated LNPs, relieving self-quenching and generating fluorescence. (d) Beacon-response analysis across increasing beacon concentrations, with free TLS11a included as a reference control to define assay performance and estimate accessible aptamer signal. Signal intensity should be plotted in fluorescence units with the linear range clearly indicated. These supplementary data support the stepwise particle assembly shown in Fig. 1 and the orthogonal aptamer-display validation presented in Fig. 2 when interpreted with the main physicochemical analyses.

**Supplementary Figure S2.**
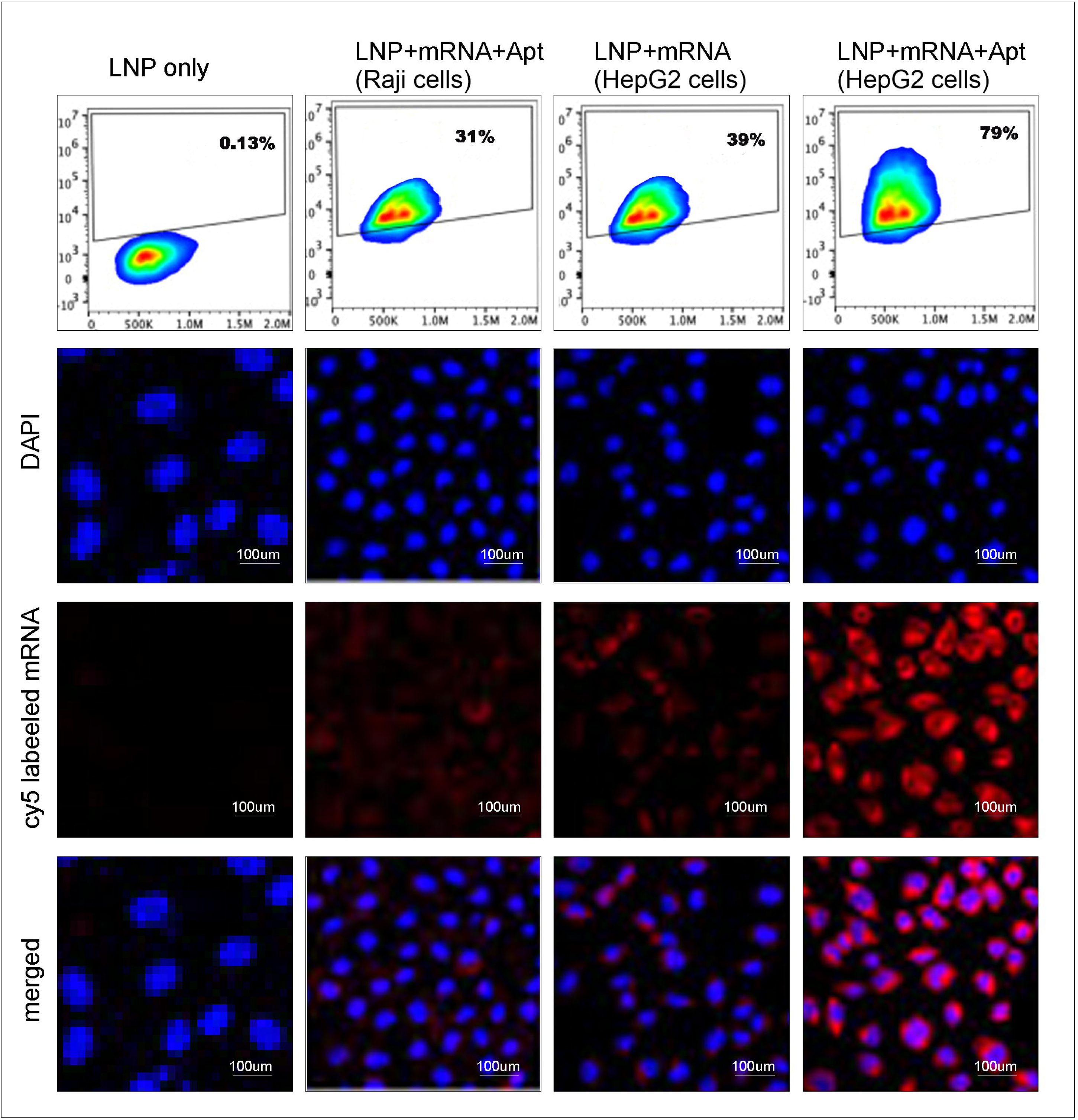
TLS11a-associated preferential uptake of Cy5-mRNA in HepG2 cells. Cy5-labeled mRNA delivery was evaluated by flow cytometry and confocal microscopy across four conditions: LNP-only control in HepG2 cells, TLS11a-functionalized cargo-loaded LNPs in Raji cells, non-functionalized cargo-loaded LNPs in HepG2 cells, and TLS11a-functionalized cargo-loaded LNPs in HepG2 cells. Flow-cytometry density plots show the Cy5-positive fractions for each condition, with minimal background signal in the LNP-only control and increased Cy5 positivity in HepG2 cells after TLS11a functionalization. Uptake was higher in TLS11a-functionalized HepG2 cells than in Raji cells exposed to the same formulation, supporting preferential HepG2-associated delivery rather than absolute cell specificity. Confocal microscopy shows DAPI-stained nuclei in blue and Cy5-mRNA signal in red, with the strongest Cy5 signal observed in TLS11a-functionalized HepG2 cells. Scale bars, 100 µm. These data support TLS11a-associated enhancement of mRNA uptake and complement the delivery analysis presented in the main text.

**Supplementary Figure S3.**
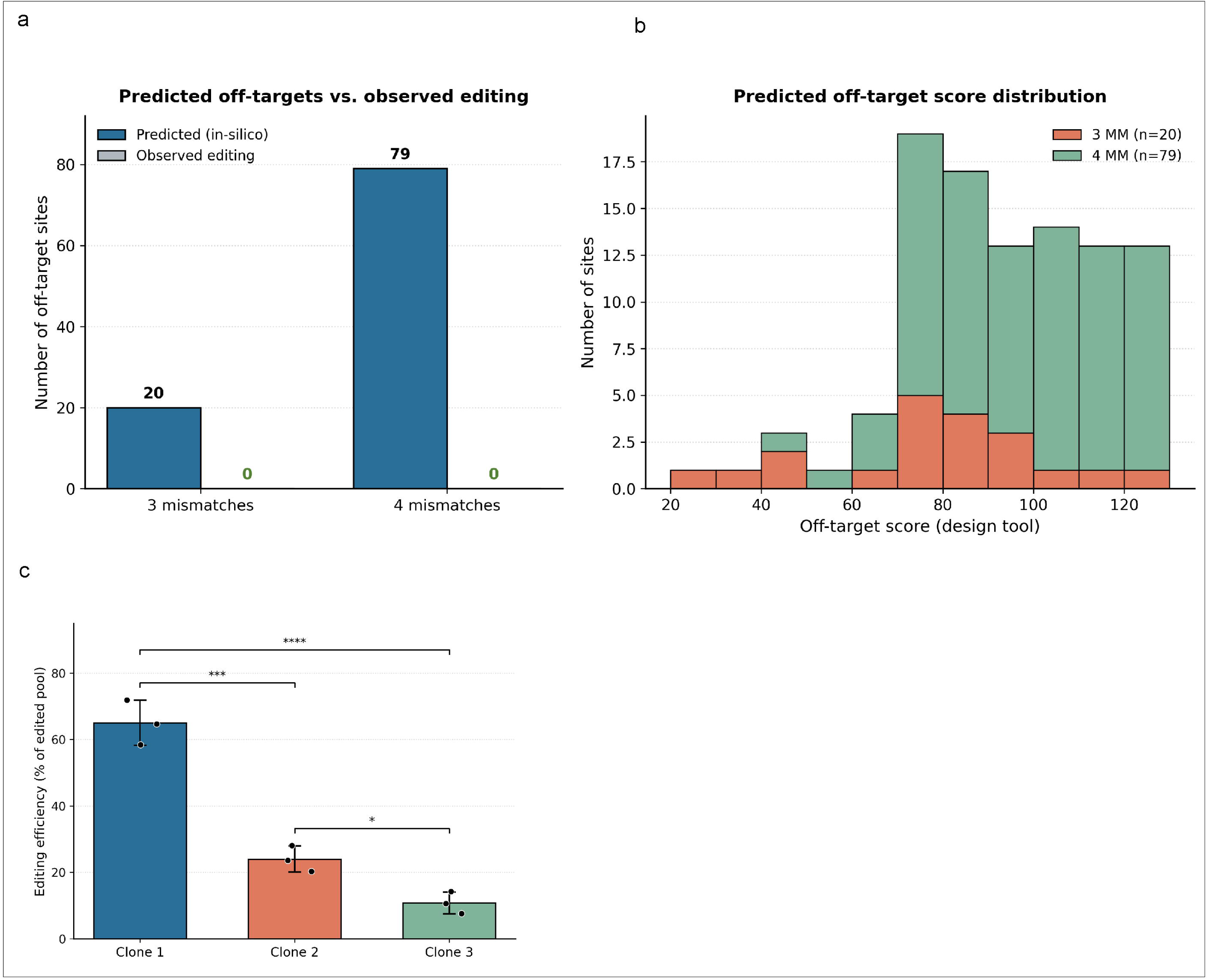
*In-silico* off-target assessment and clonal recovery after LDLR exon 2 editing. (a) Computational off-target prediction for the LDLR exon 2 sgRNA identified 20 candidate sites at 3-mismatch stringency and 79 candidate sites at 4-mismatch stringency. No editing was detected at the screened candidate loci using the assay described in Methods (PCR amplification followed by Sanger sequencing with ICE deconvolution; limit of detection approximately 5% indels), within the stated detection sensitivity. (b) Distribution of design-tool off-target scores for predicted 3-mismatch and 4-mismatch sites, illustrating the relative risk profile of candidate genomic loci (for the design-tool cutting-frequency-determination (CFD) score used here, a higher score indicates greater sequence similarity to the on-target site and therefore higher predicted off-target risk) prioritized for experimental screening. (c) Relative recovery of the three validated single-cell-derived LDLR-edited clones after expansion, shown as Clone 1, Clone 2, and Clone 3. These values describe the proportion of recovered validated clones (individual data points in panel c represent independent single-cell cloning experiments, not bulk editing efficiency) within the edited clonal pool and should not be interpreted as bulk editing efficiency. Bulk editing was measured separately by T7E1 analysis in Fig. 3e, while Sanger-confirmed indel sequences are shown in Fig. 3d. Bars show mean ± s.d.; individual data points represent independent measurements.

**Supplementary Figure S4.**
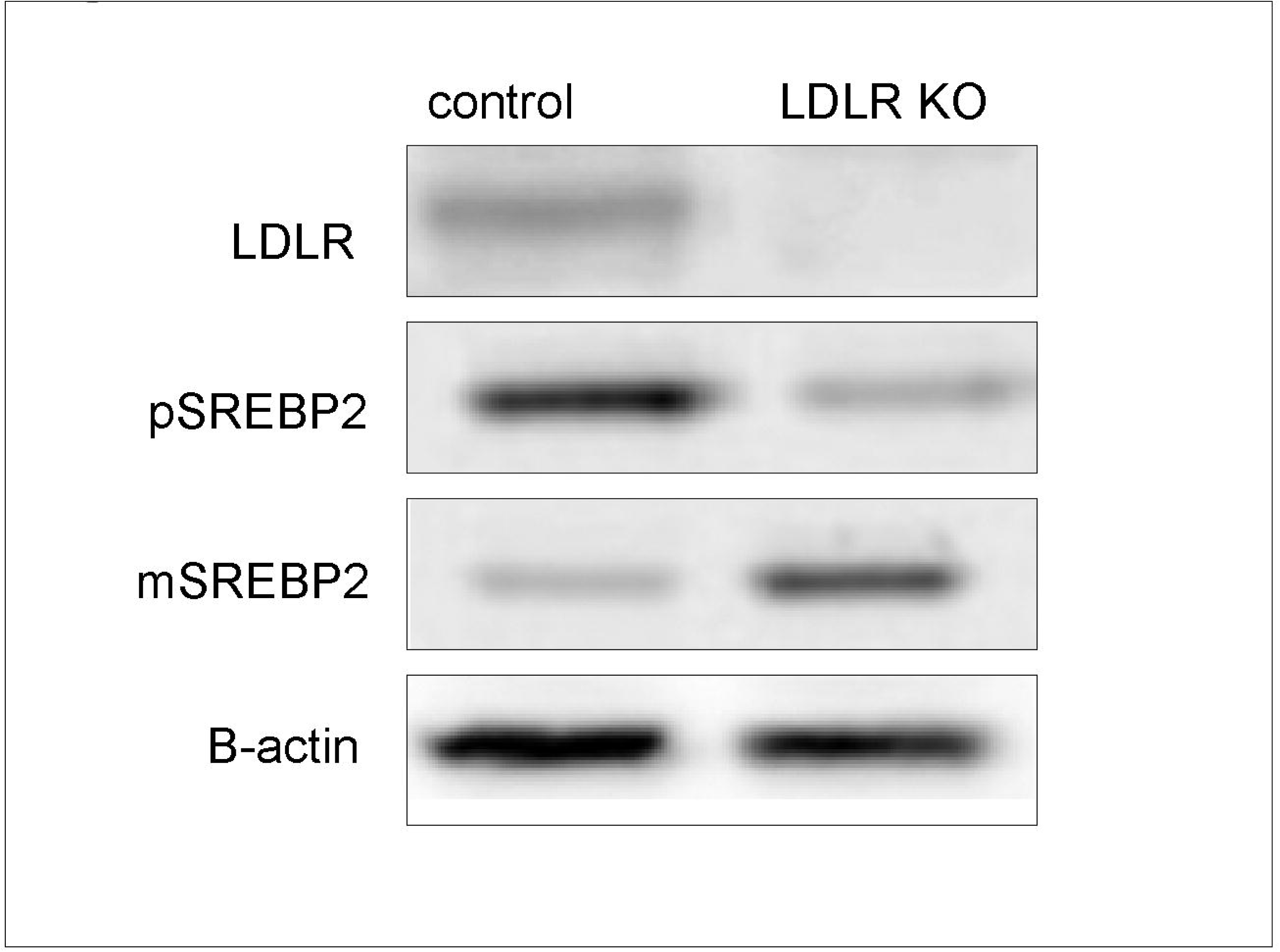
Representative immunoblot of LDLR loss and SREBP-2 processing in LDLR-KO HepG2 cells. Representative Western blot of control and LDLR-KO HepG2 cells probed for LDLR, precursor SREBP-2, mature SREBP-2, and β-actin loading control. LDLR protein was undetectable in LDLR-KO cells under the exposure conditions shown, supporting protein-level validation of LDLR disruption. LDLR-KO cells showed reduced precursor SREBP-2 signal and increased mature SREBP-2 signal, consistent with enhanced sterol-responsive SREBP-2 processing in cholesterol-depleted cells. This blot supports the densitometric quantification shown in Fig. 5i to k and is representative of three independent experiments.

## Supplementary Tables

**Supplementary Table S1.**
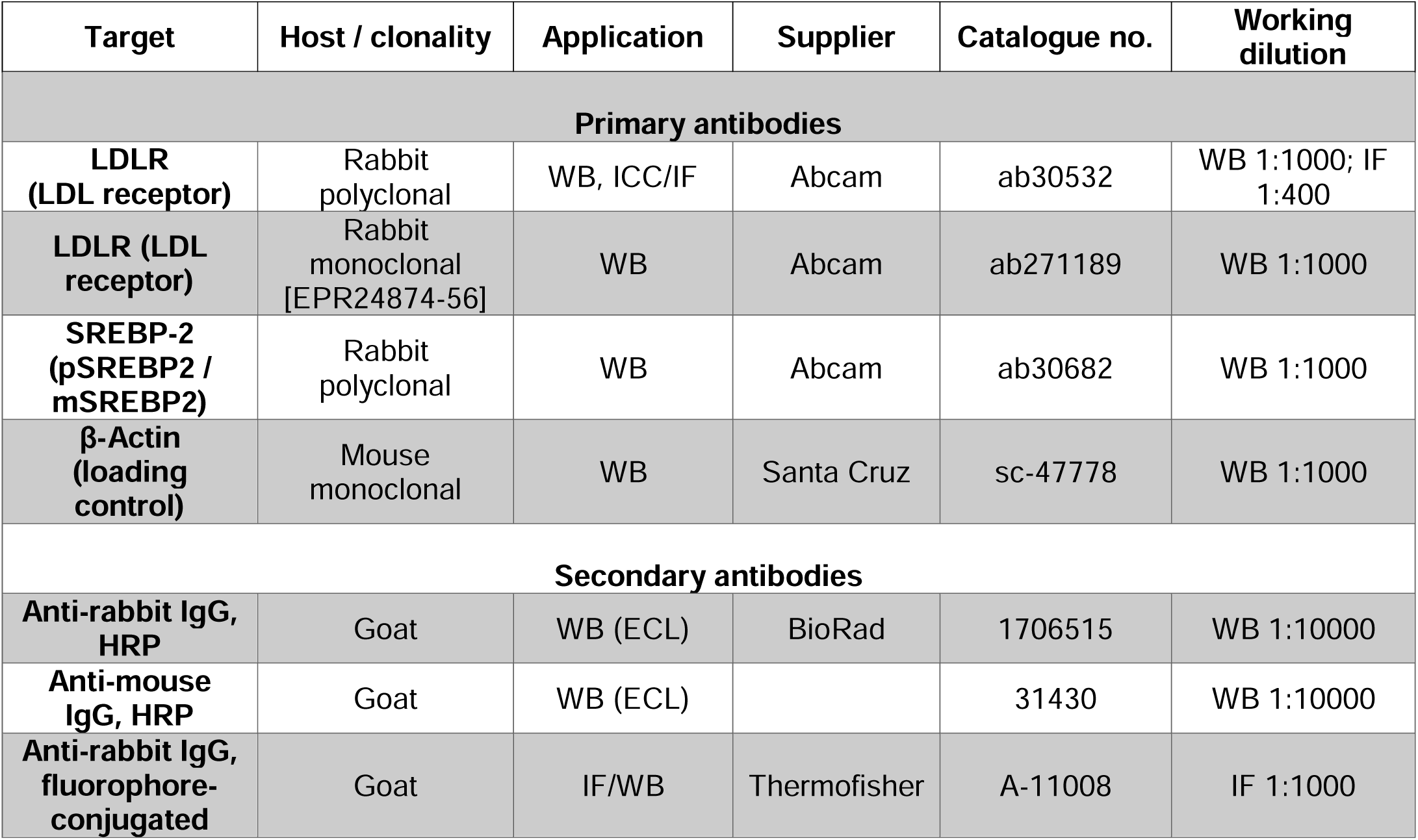
Primary and Secondary Antibodies Used in This Study.

**Supplementary Table S2.**
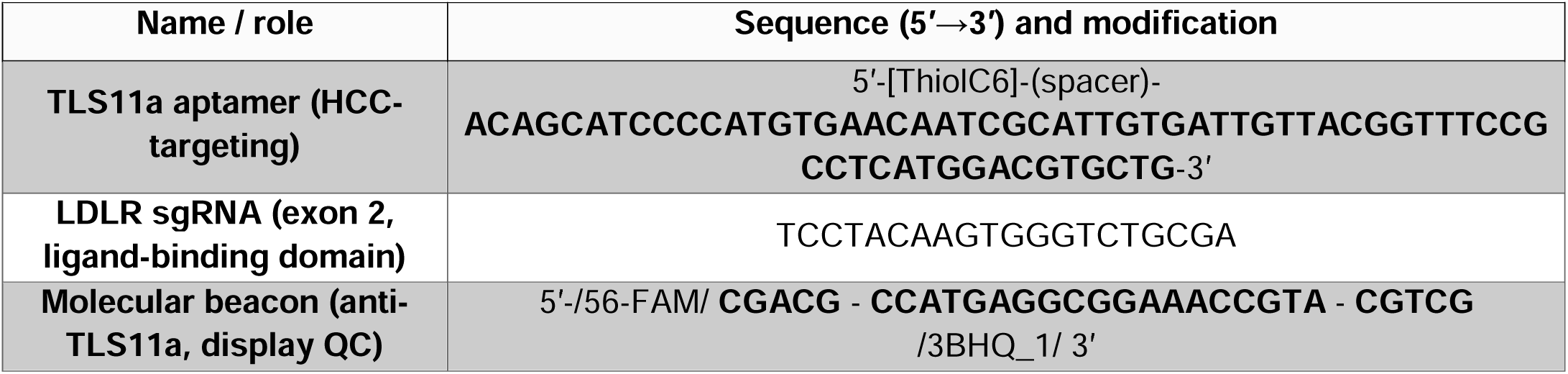
TLS11a Aptamer and LDLR-Targeting Guide RNA Sequences.

**Supplementary Table S3.**
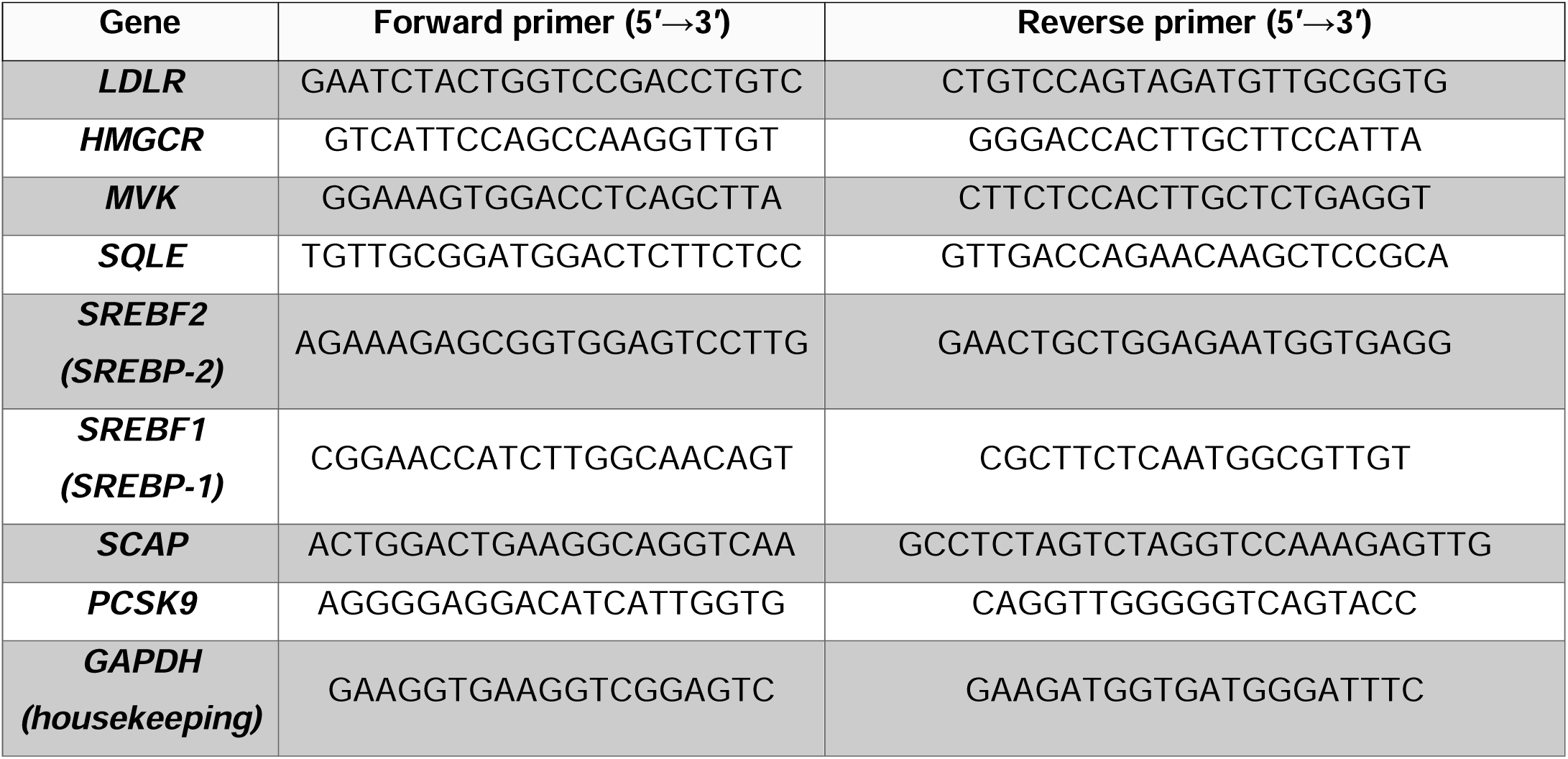
Primer sequences used for RT-qPCR analysis.

**Supplementary Table S4.**
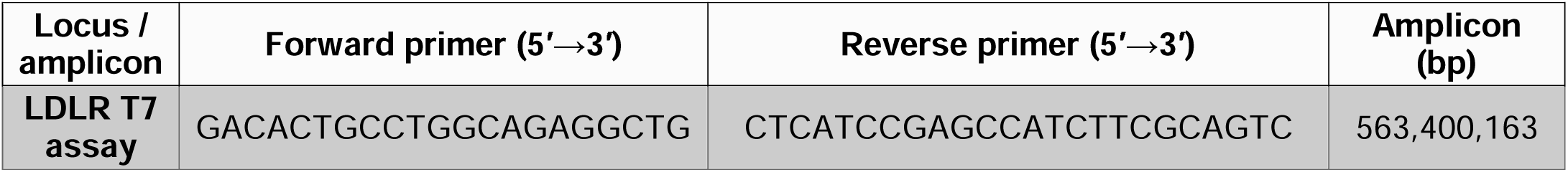
Primer Sequences for the T7 Endonuclease I (T7E1) Assay.

## Acknowledgements

The authors gratefully acknowledge the Heart Centre of Excellence and the Centre for Genomic Medicine (Department of Genomic Medicine), King Faisal Specialist Hospital & Research Centre, Riyadh, Saudi Arabia, for their support of this study. We thank these centres for providing access to the infrastructure, core facilities, and scientific expertise that made this work possible, and for their continued encouragement throughout the project.

## Author Contributions

I.H. conceived and designed the study, carried out the majority of the experimental work, analysed and interpreted the data, and wrote the original manuscript draft. R.E. performed a subset of the experiments and contributed to data analysis and manuscript writing. T.A.M. and F.R. carried out the lipid nanoparticle (LNP) formulation and aptamer conjugation and characterisation work. H.A. and J.S. performed the cell-based experiments. N.K. contributed clinical and cardiovascular expertise and interpreted the findings in the context of familial hypercholesterolemia. A.M.A. contributed to the genomic and CRISPR editing analyses and provided resources. F.S. (Farhatullah Syed) supervised the project, provided overall direction and resources, and critically revised the manuscript. All authors reviewed, edited, and approved the final version of the manuscript and agree to be accountable for all aspects of the work.

## Data Availability Statement

All data generated or analyzed during this study are included in this published article and its Supplementary Information. Additional datasets and raw data are available from the corresponding author upon reasonable request.

## Ethics Declarations

This study did not involve human participants, human-derived primary samples, or animal experimentation. All work was performed using established, commercially available human cell lines (HepG2 and Raji) and therefore did not require ethical approval.The authors confirm that any use of generative artificial-intelligence tools was limited to language editing and take full responsibility for the integrity and accuracy of the content, in line with COPE guidance on authorship and AI tools [48].

## Consent for Publication

Not applicable.

## References

1. Hou X, Zaks T, Langer R, Dong Y. Lipid nanoparticles for mRNA delivery. Nat Rev Mater. 2021;6(12):1078–1094. 10.1038/s41578-021-00358-0

2. Kulkarni JA, Witzigmann D, Chen S, Cullis PR, van der Meel R. Lipid nanoparticle technology for clinical translation of siRNA therapeutics. Acc Chem Res. 2019;52(9):2435–2444. 10.1021/acs.accounts.9b00368

3. Cullis PR, Hope MJ. Lipid nanoparticle systems for enabling gene therapies. Mol Ther. 2017;25(7):1467–1475. 10.1016/j.ymthe.2017.03.013

4. Carrasco MJ, Alishetty S, Alameh MG, Said H, Wright L, Paige M, et al. Ionization and structural properties of mRNA lipid nanoparticles influence expression in intramuscular and intravascular administration. Commun Biol. 2021;4:956. 10.1038/s42003-021-02441-2

5. Gilleron J, Querbes W, Zeigerer A, Borodovsky A, Marsico G, Schubert U, et al. Image-based analysis of lipid nanoparticle-mediated siRNA delivery, intracellular trafficking and endosomal escape. Nat Biotechnol. 2013;31(7):638–646. 10.1038/nbt.2612

6. Kauffman KJ, Dorkin JR, Yang JH, Heartlein MW, DeRosa F, Mir FF, et al. Optimization of lipid nanoparticle formulations for mRNA delivery in vivo with fractional factorial and definitive screening designs. Nano Lett. 2015;15(11):7300–7306. 10.1021/acs.nanolett.5b02497

7. Jayaraman M, Ansell SM, Mui BL, Tam YK, Chen J, Du X, et al. Maximizing the potency of siRNA lipid nanoparticles for hepatic gene silencing in vivo. Angew Chem Int Ed. 2012;51(34):8529–8533. 10.1002/anie.201203263

8. Akinc A, Querbes W, De S, Qin J, Frank-Kamenetsky M, Jayaprakash KN, et al. Targeted delivery of RNAi therapeutics with endogenous and exogenous ligand-based mechanisms. Mol Ther. 2010;18(7):1357–1364. 10.1038/mt.2010.85

9. Dilliard SA, Siegwart DJ. Passive, active and endogenous organ-targeted lipid and polymer nanoparticles for delivery of genetic drugs. Nat Rev Mater. 2023;8(4):282–300. 10.1038/s41578-022-00529-7

10. Kim M, Jeong M, Hur S, Cho Y, Park J, Jung H, et al. Engineered ionizable lipid nanoparticles for targeted delivery of RNA therapeutics into different types of cells in the liver. Sci Adv. 2021;7(9):eabf4398. 10.1126/sciadv.abf4398

11. Nag OK, Delehanty JB. Active cellular and subcellular targeting of nanoparticles for drug delivery. Pharmaceutics. 2019;11(10):543. 10.3390/pharmaceutics11100543

12. Marques AC, Costa PJ, Velho S, Amaral MH. Functionalizing nanoparticles with cancer-targeting antibodies: a comparison of strategies. J Control Release. 2020;320:180–200. 10.1016/j.jconrel.2020.01.035

13. Zhou J, Rossi J. Aptamers as targeted therapeutics: current potential and challenges. Nat Rev Drug Discov. 2017;16(3):181–202. 10.1038/nrd.2016.199

14. Keefe AD, Pai S, Ellington A. Aptamers as therapeutics. Nat Rev Drug Discov. 2010;9(7):537–550. 10.1038/nrd3141

15. Meng HM, Liu H, Kuai H, Peng R, Mo L, Zhang XB. Aptamer-integrated DNA nanostructures for biosensing, bioimaging and cancer therapy. Chem Soc Rev. 2016;45(9):2583–2602. 10.1039/C5CS00645G

16. Fu Z, Xiang J. Aptamer-functionalized nanoparticles in targeted delivery and cancer therapy. Int J Mol Sci. 2020;21(23):9123. 10.3390/ijms21239123

17. Farokhzad OC, Cheng J, Teply BA, Sherifi I, Jon S, Kantoff PW, et al. Targeted nanoparticle-aptamer bioconjugates for cancer chemotherapy in vivo. Proc Natl Acad Sci USA. 2006;103(16):6315–6320. 10.1073/pnas.0601755103

18. Shangguan D, Meng L, Cao ZC, Xiao Z, Fang X, Li Y, et al. Identification of liver cancer-specific aptamers using whole live cells. Anal Chem. 2008;80(3):721–728. 10.1021/ac701962v

19. Bing T, Shangguan D, Wang Y. Facile discovery of cell-selective and target-specific aptamers by live cell-based SELEX. Mol Ther Nucleic Acids. 2015;4:e260. 10.1038/mtna.2015.32

20. Nordestgaard BG, Chapman MJ, Humphries SE, Ginsberg HN, Masana L, Descamps OS, et al. Familial hypercholesterolaemia is underdiagnosed and undertreated in the general population: guidance for clinicians to prevent coronary heart disease. Eur Heart J. 2013;34(45):3478–3490. 10.1093/eurheartj/eht273

21. Defesche JC, Gidding SS, Harada-Shiba M, Hegele RA, Santos RD, Wierzbicki AS. Familial hypercholesterolaemia. Nat Rev Dis Primers. 2017;3:17093. 10.1038/nrdp.2017.93

22. Cuchel M, Bruckert E, Ginsberg HN, Raal FJ, Santos RD, Hegele RA, et al. Homozygous familial hypercholesterolaemia: new insights and guidance for clinicians to improve detection and clinical management. Eur Heart J. 2014;35(32):2146–2157. 10.1093/eurheartj/ehu274

23. Beheshti SO, Madsen CM, Varbo A, Nordestgaard BG. Worldwide prevalence of familial hypercholesterolemia: meta-analyses of 11 million subjects. J Am Coll Cardiol. 2020;75(20):2553–2566. 10.1016/j.jacc.2020.03.057

24. Brown MS, Goldstein JL. A receptor-mediated pathway for cholesterol homeostasis. Science. 1986;232(4746):34–47. 10.1126/science.3513311

25. Goldstein JL, Brown MS. The LDL receptor. Arterioscler Thromb Vasc Biol. 2009;29(4):431–438. 10.1161/ATVBAHA.108.179564

26. Brown MS, Goldstein JL. A proteolytic pathway that controls the cholesterol content of membranes, cells, and blood. Proc Natl Acad Sci USA. 1999;96(20):11041–11048. 10.1073/pnas.96.20.11041

27. Horton JD, Goldstein JL, Brown MS. SREBPs: activators of the complete program of cholesterol and fatty acid synthesis in the liver. J Clin Invest. 2002;109(9):1125–1131. 10.1172/JCI15593

28. Brown MS, Radhakrishnan A, Goldstein JL. Retrospective on cholesterol homeostasis: the central role of Scap. Annu Rev Biochem. 2018;87:783–807. 10.1146/annurev-biochem-062917-011852

29. Horton JD, Cohen JC, Hobbs HH. PCSK9: a convertase that coordinates LDL catabolism. J Lipid Res. 2009;50(Suppl):S172–S177. 10.1194/jlr.R800091-JLR200

30. Rong S, Cortés VA, Rashid S, Anderson NN, McDonald JG, Liang G, et al. Expression of SREBP-1c requires SREBP-2-mediated generation of a sterol ligand for LXR in livers of mice. eLife. 2017;6:e25015. 10.7554/eLife.25015

31. Jinek M, Chylinski K, Fonfara I, Hauer M, Doudna JA, Charpentier E. A programmable dual-RNA-guided DNA endonuclease in adaptive bacterial immunity. Science. 2012;337(6096):816–821. 10.1126/science.1225829

32. Cong L, Ran FA, Cox D, Lin S, Barretto R, Habib N, et al. Multiplex genome engineering using CRISPR/Cas systems. Science. 2013;339(6121):819–823. 10.1126/science.1231143

33. Doudna JA, Charpentier E. The new frontier of genome engineering with CRISPR-Cas9. Science. 2014;346(6213):1258096. 10.1126/science.1258096

34. Musunuru K, Chadwick AC, Mizoguchi T, Garcia SP, DeNizio JE, Reiss CW, et al. In vivo CRISPR base editing of PCSK9 durably lowers cholesterol in primates. Nature. 2021;593(7859):429–434. 10.1038/s41586-021-03534-y

35. Lee RG, Mazzola AM, Braun MC, Platt C, Vafai SB, Kathiresan S, et al. Efficacy and safety of an investigational single-course CRISPR base-editing therapy targeting PCSK9 in nonhuman primate and mouse models. Circulation. 2023;147(3):242–253. 10.1161/CIRCULATIONAHA.122.062132

36. Rosenblum D, Gutkin A, Kedmi R, Ramishetti S, Veiga N, Jacobi AM, et al. CRISPR-Cas9 genome editing using targeted lipid nanoparticles for cancer therapy. Sci Adv. 2020;6(47):eabc9450. 10.1126/sciadv.abc9450

37. Xing H, Tang L, Yang X, Hwang K, Wang W, Yin Q, et al. Selective delivery of an anticancer drug with aptamer-functionalized liposomes to breast cancer cells in vitro and in vivo. J Mater Chem B. 2013;1(39):5288–5297. 10.1039/c3tb20412j

38. Kypr J, Kejnovská I, Renčiuk D, Vorlíčková M. Circular dichroism and conformational polymorphism of DNA. Nucleic Acids Res. 2009;37(6):1713–1725. 10.1093/nar/gkp026

39. Baer DR, Engelhard MH. XPS analysis of nanostructured materials and biological surfaces. J Electron Spectrosc Relat Phenom. 2010;178–179:415–432. 10.1016/j.elspec.2009.09.003

40. Tyagi S, Kramer FR. Molecular beacons: probes that fluoresce upon hybridization. Nat Biotechnol. 1996;14(3):303–308. 10.1038/nbt0396-303

41. Ran FA, Hsu PD, Wright J, Agarwala V, Scott DA, Zhang F. Genome engineering using the CRISPR-Cas9 system. Nat Protoc. 2013;8(11):2281–2308. 10.1038/nprot.2013.143

42. Doench JG, Fusi N, Sullender M, Hegde M, Vaimberg EW, Donovan KF, et al. Optimized sgRNA design to maximize activity and minimize off-target effects of CRISPR-Cas9. Nat Biotechnol. 2016;34(2):184–191. 10.1038/nbt.3437

43. Guschin DY, Waite AJ, Katibah GE, Miller JC, Holmes MC, Rebar EJ. A rapid and general assay for monitoring endogenous gene modification. Methods Mol Biol. 2010;649:247–256. 10.1007/978-1-60761-753-2_15

44. Sentmanat MF, Peters ST, Florian CP, Connelly JP, Pruett-Miller SM. A survey of validation strategies for CRISPR-Cas9 editing. Sci Rep. 2018;8:888. 10.1038/s41598-018-19441-8

45. Livak KJ, Schmittgen TD. Analysis of relative gene expression data using real-time quantitative PCR and the 2^−ΔΔCt method. Methods. 2001;25(4):402–408. 10.1006/meth.2001.1262

46. Goldstein JL, Basu SK, Brown MS. Receptor-mediated endocytosis of low-density lipoprotein in cultured cells. Methods Enzymol. 1983;98:241–260. 10.1016/0076-6879(83)98152-1

47. Maxfield FR, Wüstner D. Analysis of cholesterol trafficking with fluorescent probes. Methods Cell Biol. 2012;108:367–393. 10.1016/B978-0-12-386487-1.00017-1

48. Committee on Publication Ethics (COPE). Authorship and AI tools: COPE position statement. 2023. 10.24318/cCVRZBms

49. Semple SC, Akinc A, Chen J, Sandhu AP, Mui BL, Cho CK, et al. Rational design of cationic lipids for siRNA delivery. Nat Biotechnol. 2010;28(2):172–176. 10.1038/nbt.1602

50. MacArthur JM, Bishop JR, Stanford KI, Wang L, Bensadoun A, Witztum JL, et al. Liver heparan sulfate proteoglycans mediate clearance of triglyceride-rich lipoproteins independently of LDL receptor family members. J Clin Invest. 2007;117(1):153–164. 10.1172/JCI29154

51. Rohlmann A, Gotthardt M, Hammer RE, Herz J. Inducible inactivation of hepatic LRP gene by Cre-mediated recombination confirms role of LRP in clearance of chylomicron remnants. J Clin Invest. 1998;101(3):689–695. 10.1172/JCI1240

52. Acton S, Rigotti A, Landschulz KT, Xu S, Hobbs HH, Krieger M. Identification of scavenger receptor SR-BI as a high density lipoprotein receptor. Science. 1996;271(5248):518–520. 10.1126/science.271.5248.518

53. Lucero D, Dikilitas O, Mendelson MM, Aligabi Z, Islam P, Neufeld EB, Bansal AT, Freeman LA, Vaisman B, Tang J, Combs CA, Li Y, Voros S, Kullo IJ, Remaley AT. Transgelin: a new gene involved in LDL endocytosis identified by a genome-wide CRISPR-Cas9 screen. J Lipid Res. 2022 Jan;63(1):100160. doi: 10.1016/j.jlr.2021.100160. Epub 2021 Dec 10. PMID: 34902367; PMCID: PMC8953622.

