## Supplementary Tables for "A TLS11a-decorated ionizable lipid nanoparticle platform and a multilevel-validated CRISPR LDLR-knockout HepG2 model for hepatocyte-preferential mRNA delivery"

**Supplementary Table S1. Primary and Secondary Antibodies Used in This Study**

| Target | Host / clonality | Application | Supplier | Catalogue no. | Working dilution |
| --- | --- | --- | --- | --- | --- |
| Primary antibodies | | | | | |
| LDLR  (LDL receptor) | Rabbit polyclonal | WB, ICC/IF | Abcam | ab30532 | WB 1:1000; IF 1:400 |
| LDLR (LDL receptor) | Rabbit monoclonal [EPR24874-56] | WB | Abcam | ab271189 | WB 1:1000 |
| SREBP-2 (pSREBP2 / mSREBP2) | Rabbit polyclonal | WB | Abcam | ab30682 | WB 1:1000 |
| β-Actin (loading control) | Mouse monoclonal | WB | Santa Cruz | sc-47778 | WB 1:1000 |
| Secondary antibodies | | | | | |
| Anti-rabbit IgG, HRP | Goat | WB (ECL) | BioRad | 1706515 | WB 1:10000 |
| Anti-mouse IgG, HRP | Goat | WB (ECL) |  | 31430 | WB 1:10000 |
| Anti-rabbit IgG, fluorophore-conjugated | Goat | IF/WB | Thermofisher | A-11008 | IF 1:1000 |

**Supplementary Table S2. TLS11a Aptamer and LDLR-Targeting Guide RNA Sequences**

| Name / role | Sequence (5′→3′) and modification |
| --- | --- |
| TLS11a aptamer (HCC-targeting) | 5′-[ThiolC6]-(spacer)-**ACAGCATCCCCATGTGAACAATCGCATTGTGATTGTTACGGTTTCCG**  **CCTCATGGACGTGCTG**-3′ |
| LDLR sgRNA (exon 2, ligand-binding domain) | TCCTACAAGTGGGTCTGCGA |
| Molecular beacon (anti-TLS11a, display QC) | 5′-/56-FAM/ **CGACG** - **CCATGAGGCGGAAACCGTA** - **CGTCG** /3BHQ_1/ 3′ |

**Supplementary Table S3. Primer sequences used for RT-qPCR analysis.**

| Gene | Forward primer (5′→3′) | Reverse primer (5′→3′) |
| --- | --- | --- |
| *LDLR* | GAATCTACTGGTCCGACCTGTC | CTGTCCAGTAGATGTTGCGGTG |
| *HMGCR* | GTCATTCCAGCCAAGGTTGT | GGGACCACTTGCTTCCATTA |
| *MVK* | GGAAAGTGGACCTCAGCTTA | CTTCTCCACTTGCTCTGAGGT |
| *SQLE* | TGTTGCGGATGGACTCTTCTCC | GTTGACCAGAACAAGCTCCGCA |
| *SREBF2 (SREBP-2)* | AGAAAGAGCGGTGGAGTCCTTG | GAACTGCTGGAGAATGGTGAGG |
| *SREBF1 (SREBP-1)* | CGGAACCATCTTGGCAACAGT | CGCTTCTCAATGGCGTTGT |
| *SCAP* | ACTGGACTGAAGGCAGGTCAA | GCCTCTAGTCTAGGTCCAAAGAGTTG |
| *PCSK9* | AGGGGAGGACATCATTGGTG | CAGGTTGGGGGTCAGTACC |
| *GAPDH (housekeeping)* | GAAGGTGAAGGTCGGAGTC | GAAGATGGTGATGGGATTTC |

**Supplementary Table S4. Primer Sequences for the T7 Endonuclease I (T7E1) Assay**

| Locus / amplicon | Forward primer (5′→3′) | Reverse primer (5′→3′) | Amplicon (bp) |
| --- | --- | --- | --- |
| LDLR T7 assay | GACACTGCCTGGCAGAGGCTG | CTCATCCGAGCCATCTTCGCAGTC | 563,400,163 |
